# Effects of Age and Diet on Triglyceride Metabolism in Mice

**DOI:** 10.1101/2024.07.19.602944

**Authors:** Kathryn M. Spitler, Shwetha K. Shetty, Brandon S.J. Davies

## Abstract

**Background:** Both age and diet can contribute to alterations in triglyceride metabolism and subsequent metabolic disease. In humans, plasma triglyceride levels increase with age. Diets high in saturated fats can increase triglyceride levels while diets high in omega-3 fatty acids decrease triglyceride levels. Here we asked how age and long-term diet effected triglyceride metabolism in mice.

**Methods:** We fed male and female mice a low-fat diet, a western diet, or a diet high in polyunsaturated and omega-3 (n-3) fatty acids for up to 2 years. We measured survival, body composition, plasma triglyceride levels, chylomicron clearance, and oral fat, glucose, and insulin tolerance.

**Results:** Triglyceride levels in mice did not increase with age, regardless of diet. Oral fat tolerance increased with age, while chylomicron clearance remained unchanged. Mice fed western diet had decreased survival. Interestingly, mice fed the n-3 diet gained more lean mass, and had lower insulin levels than mice fed either low-fat or western diet. Moreover, triglyceride uptake into the hearts of mice fed the n-3 diet was strikingly higher than in other groups.

**Conclusions:** In mice, age-induced changes in triglyceride metabolism did not match those in humans. Our data suggested that mice, like humans, had decreased fat absorption with age, but plasma triglyceride clearance did not decrease with age in mice, resulting in lower plasma triglyceride levels and improved oral fat tolerance with age. A chronic diet high in n-3 fatty acids increased insulin sensitivity and uptake of triglycerides specifically into the heart but how these observations are connected is unclear.

**Research Perspectives:**

- The changes in triglyceride metabolism that occur with age in humans are not reflected in a mouse model, thus mice are likely not an ideal model for understanding how age impacts lipid metabolism and subsequent metabolic disease.
- A fish-oil based high-fat diet high in omega-3 fatty acids significantly increases fatty acid uptake in the heart while at the same time decreases fasting insulin levels.
- In future studies it will be important to understand how the omega-3 fatty acid induced increase in fatty acid uptake affects cardiac function and how it is related to other phenotypes induced by omega-3 fatty acids.

## Introduction

The world population is seeing an increase in both age and obesity. In the United States, more than 20% of the population will be more than 65 years of age by the year 2030.^1^ Increasing evidence has shown that during aging changes in body weight, fat accumulation, insulin resistance and triglyceride metabolism are associated with the development of chronic metabolic diseases, such as obesity, type 2 diabetes and hyperlipidemia.^2–4^ The use of animal models to model human diseases is the mainstay of scientific studies that aim to elucidate the molecular mechanisms that mediate disease. Currently, data demonstrating that mice can be used as a reliable model for obesity and metabolic disease during aging is lacking.

The consumption of high-fat food is a contributing factor to the development of metabolic diseases.^5^ In aging adults the consumption of high fat foods has increased, in part due to convenience, modern lifestyles, and overall palatability of these foods.^6^ The long-term consequences of a high-fat diet can be gradual and cumulative and intertwine with other factors such as age and obesity.^7^ The hallmark of obesogenic studies in mice has been the utilization of high fat diets. However, the composition of these diets can vary across many parameters, including the source of fat, the overall percentage of fat (anywhere from 40-60% of kcal), and the addition of additional components such as cholesterol.^8^ The vast majority of studies have utilized these diets for relatively short-term studies: 12 weeks to 6 months. How diets alter metabolic phenotypes of mice over their life-time (average lifespan of the laboratory mouse is 2-2.5 years) is not fully understood.

In contrast to diets high in saturated fatty acids, increased consumption of omega-3 (n-3) fatty acids has been proposed as a way to mitigate metabolic disease. Outcomes from the JELIS study in Japan and the Reduce-IT study, which mostly studied Caucasians, showed that high doses of the omega-3 fatty acid eicosapentaenoic acid (EPA) reduced cardiovascular risk in humans and lowered plasma triglyceride levels,^9,10^ but the mechanism by which this occurs is not known. Other human studies have shown that increased consumption of omega-3 fatty acids reduces fasting VLDL, fasting plasma triglyceride (TG) levels, and postprandial TG-rich lipoprotein levels.^11–13^ In rodent models, omega-3 fatty acids have been shown to have many potentially beneficial effects on lipid and glucose metabolism (reviewed in ^14^). However, most studies in model organisms have been done in young rodents fed test diets for a limited amount of time (16 weeks or less). There is a need to examine the effects of a chronic diet in aged models.

Animal models of metabolic disease can provide valuable insight into how metabolic homeostasis can be altered by conditions such as long-term administration of a high-fat diet.^15^ For example, C57BL/6 mice fed a high-fat diet are classically used in the literature as models of increased fasting plasma glucose levels and insulin resistance.^16^ An important factor in the utilization of animal models in biomedical research is that they accurately model human pathologies.^17^ Understanding the effects of long-term high-fat diet consumption on triglyceride metabolism during the lifespan of the mouse could provide insights into the underlying mechanisms that promote metabolic disease if mice accurately reflect the human condition. However, differences between humans and mice exist and it is important to identify and address these differences. For example, in both mice and humans weight and body fat peak between mid and late life.^18^ In contrast, glucose levels remained relatively stable in mice until late in life when levels decline, whereas, in humans glucose levels tend to increase throughout the course of life.^18^ Although several studies have observed how triglyceride metabolism is altered with age and chronic diet in humans,^19–24^ there is a paucity of information regarding the utility of mice as an appropriate model of age-driven triglyceride metabolic disturbances. The aim of this study was to determine how age, in combination with a low fat, high saturated fat, or high omega-3 fat diet alters triglyceride homeostasis in male and female C57BL/6 mice.

## Methods

### Mice

6-week-old C57BL/6 mice (Strain# 000664) were purchased from Jackson Laboratories. Following arrival at the University of Iowa, mice were allowed to acclimate for two weeks before initiation of study. At 8 weeks of age, mice were randomly assigned (using randomizer.org) to a low-fat diet (Teklad Diets TD.05230), a Western diet (Teklad Diets TD.200494), or a diet high in poly-unsaturated and omega-3 fatty acids (Teklad Diets TD.200495)(**Table 1**). 8 mice of each gender were assigned to each cohort, with the exception of the female, 2-year cohorts. For these cohorts 10 mice were used. A total of 150 mice were used in this study. Sample size was calculated using power analysis as to have 80% power to detect 40% differences with a p value of 0.05 for triglyceride uptake assays. Researchers were not blinded to diet assignment. Potential confounders such as order of measurements between diet cohorts were not controlled. Mice were housed 4 per cage. All animals were maintained at 25°C with a 12:12 hour light-dark cycle. Water was provided *ad libitum*. Mice were fed their respective diets until the time of sacrifice (age of 16 weeks, 1 year, or 2 years). Mice were weighed weekly. For food consumption studies food from each cage was weighed daily at the same time for one week. Survival rate was tracked for mice in the two-year cohort. Death included natural death as well as humane euthanasia due to severe health issues including significant weight loss, untreatable wounds or sickness, or inability to rise or ambulate.

**Table 1:** Diets used in this study. LFD=low fat diet. WD=Western Diet. n3D= omega-3 enriched diet.

|  | **LFD** | **WD** | **n3D** |
| --- | --- | --- | --- |
| **Protein, % by weight** | 17.3 | 17.3 | 17.3 |
| **Protein, % kcal from** | 18.7 | 15.2 | 15.2 |
| **Carbohydrate, % by weight** | 63.5 | 48.5 | 48.9 |
| **Carbohydrate, % kcal from** | 68.7 | 42.7 | 43.0 |
| **Sucrose, g/kg** | 341 | 341.5 | 341.5 |
| **Fat, % by weight** | 5.2 | 21.2 | 21.2 |
| **Fat, % kcal from** | 12.6 | 42 | 41.9 |
| **Saturated fat g/kg** | 25.4 | 132.3 | 57.0 |
| **Monounsaturated fat g/kg** | 13.7 | 60.9 | 46.7 |
| **Polyunsaturated fat g/kg** | 9.3 | 8.4 | 73.2 |
| **Omega-3 fat g/kg** | 1.2 | 1.1 | 60.6 |
| **n-6:n-3 ratio** | 6.8 | 8 | 0.2 |

All animal procedures were approved by the Institutional Animal Care and Use Committee at the University of Iowa and were carried out in accordance with the National Institute of Health *Guide for Care and Use of Laboratory Animals*.

### Body composition analysis

Lean muscle and fat mass was determined using nuclear magnetic resonance (NMR) in mice before the start of diet (8 weeks of age), after 8 weeks on diet (16 weeks of age), and at 8 months, 12 months, 18 months, and 24 months of age. Mice were weighed and placed into a tube restraint without anesthesia. Mice were scanned with either a Bruker LF50 (mice weighing less than 50 g) or a Bruker LF90 (mice weigher over 50 grams). Following NMR scanning mice were immediately returned to their home cages. NMR measurements were performed in the Fraternal Order of the Eagles Diabetes Research Center Metabolic Phenotyping Core at the University of Iowa.

### Fasting plasma triglyceride measurements

Fasting plasma was collected in fasted (6 h) mice at the ages indicated in the figures. Blood was collected into EDTA-coated capillary tubes and plasma was collected following centrifugation (1,500xg, 15 min, 4°C). Plasma triglycerides were analyzed in duplicate using the Infinity Triglyceride Reagent (Thermo Scientific) as previously described.^25^

### Oral Fat Tolerance Test

Oral fat tolerance tests (FTTs) were performed in mice at 13, 49, and 75 weeks, and 2 years of age after a 12 h fast. Blood was collected into an EDTA-coated capillary tube before and 1, 2, 3, 4, and 6 h after being gavaged with olive oil (10 µl/g body weight). Plasma was collected following centrifugation (1,500xg, 15 min, 4°C). Plasma was stored at −80°C until all groups had been collected. Plasma triglyceride levels were measured as above using the Infinity Triglyceride Reagent (Thermo).

### Triglyceride clearance and uptake assay

^3^H-labeled chylomicrons were prepared from *Gpihbp1^−/−^* mice as previously described.^26^ Briefly, *Gpihbp1^−/−^* mice were orally gavaged with 100 μCi of [9,10-3H(N)]-Triolein (Perkin Elmer, NET431001MC) suspended in olive oil. After 4 h, blood was collected via cardiac puncture and placed into a tube containing 0.5 M EDTA and placed on ice. The blood was then centrifuged at 1,500×g for 15 min and plasma was collected. The plasma was then centrifuged at 424,000×g twice for 2 hours at 10°C. The chylomicrons were collected from the upper layer of the resulting supernatant, resuspended in sterile saline and stored at 4°C. Radioactivity was measured using a Perkin Elmer Liquid Scintillation Counter.

Triglyceride clearance and uptake assays were performed in mice as previously described.^26^ At the end of the experimental timepoints (16 weeks, 1 year, or 2 years of age) female and male mice were fasted (6 h), anesthetized with isoflurane, and injected retro-orbitally with ^3^H-labeled chylomicrons (100 µl). Blood was collected 1, 5, 10, and 15 minutes following injection and assayed for radioactivity in BioSafe II scintillation fluid (10 µl plasma/time point). 15 minutes post injection mice were anesthetized with isoflurane and perfused with 20 ml of PBS solution containing 0.5% Tyloxapol. Tissues (heart, liver, kidney, quadricep, gonadal white adipose tissue, subcutaneous white adipose tissue, and brown adipose tissue) were harvested and weighed. Chloroform:methanol (2:1) lipid extraction was performed on 40-90 mg of each tissue. After overnight incubation at 4°C and following phase separation with 1 ml of 2M CaCl_2_ the organic and aqueous phases were assayed for radioactivity using a Beckman-Coulter Scintillation Counter. The counts per million from each fraction were combined to give total uptake CPMs. Radiolabel CPM values were normalized to the CPM of the injected dose (measured by assaying 10% of the ^3^H-Chylomicron suspension injected into the mouse.).

### Glucose and Insulin Tolerance Tests

Glucose tolerance tests (GTTs) were performed in fasted (6 h) mice at 14, 50, and 76 weeks, and 2 years of age. Blood glucose was measured with a glucometer (OneTouch Ultra) before intraperitoneal injection of glucose (1g/kg) and then 30, 60, 90 and 120 minutes post injection. At 0 and 30 minutes blood was also collected into EDTA coated capillary tubes by tail vain puncture. Blood was then centrifuged at 1500 x g for 20 min at 4°C and plasma was collected and stored at −80°C. After all blood collection, plasma insulin levels were determined using the Ultra-Sensitive Mouse Insulin ELISA Kit (Crystal Chem, 90080), according to manufacturer’s protocols.

Insulin tolerance tests (ITTs) were performed in fasted (4 h) mice at 15, 51, and 77 weeks, and 2 years of age. Blood glucose was measured with a glucometer before intraperitoneal injection of Insulin (0.5U/kg, Humalin-R) and 15, 30, 45, 60, and 90 minutes post injection.

### Statistics

Statistics were performed using Graphpad Prism 10. As decided a priori, we tested for outliers for each data set using the ROUT method with Q=1%. Outliers identified in this way were removed from graphs and from statistical analysis. The total number of mice and the number of mice excluded from individual analyses are noted in supplementary table 1. Statistical tests used for each data set are listed in the respective figure legend and in supplementary table 1. Complete lists of p values are shown in supplementary table 1.

## Results

### Experimental Design

The experimental design for this study is shown in **Figure 1A**. After 8 weeks on normal chow, mice were randomly assigned one of three diets (**Table 1**), a low-fat control diet (LFD; 12% kcal derived from fat), or one of two high-fat diets. The first high-fat diet was a classic “Western” diet (WD) with 42% of kcal from fat and a majority of the fat being milk fat-derived saturated fat. The second high-fat diet (omega-3 enriched high fat diet; n3D) also had 42% of kcal from fat, but the majority of the fat was derived from fish oil and was high in polyunsaturated and omega-3 fatty acids. For each diet, mice were assigned one of three age cohorts. The young cohort was fed their respective diet for 8 weeks. The middle-age cohort were fed their respective diets until they were 1 year old, and the old cohort until they were 2 years old. Each cohort started with 8 male and 8 female mice.

**Figure 1:**
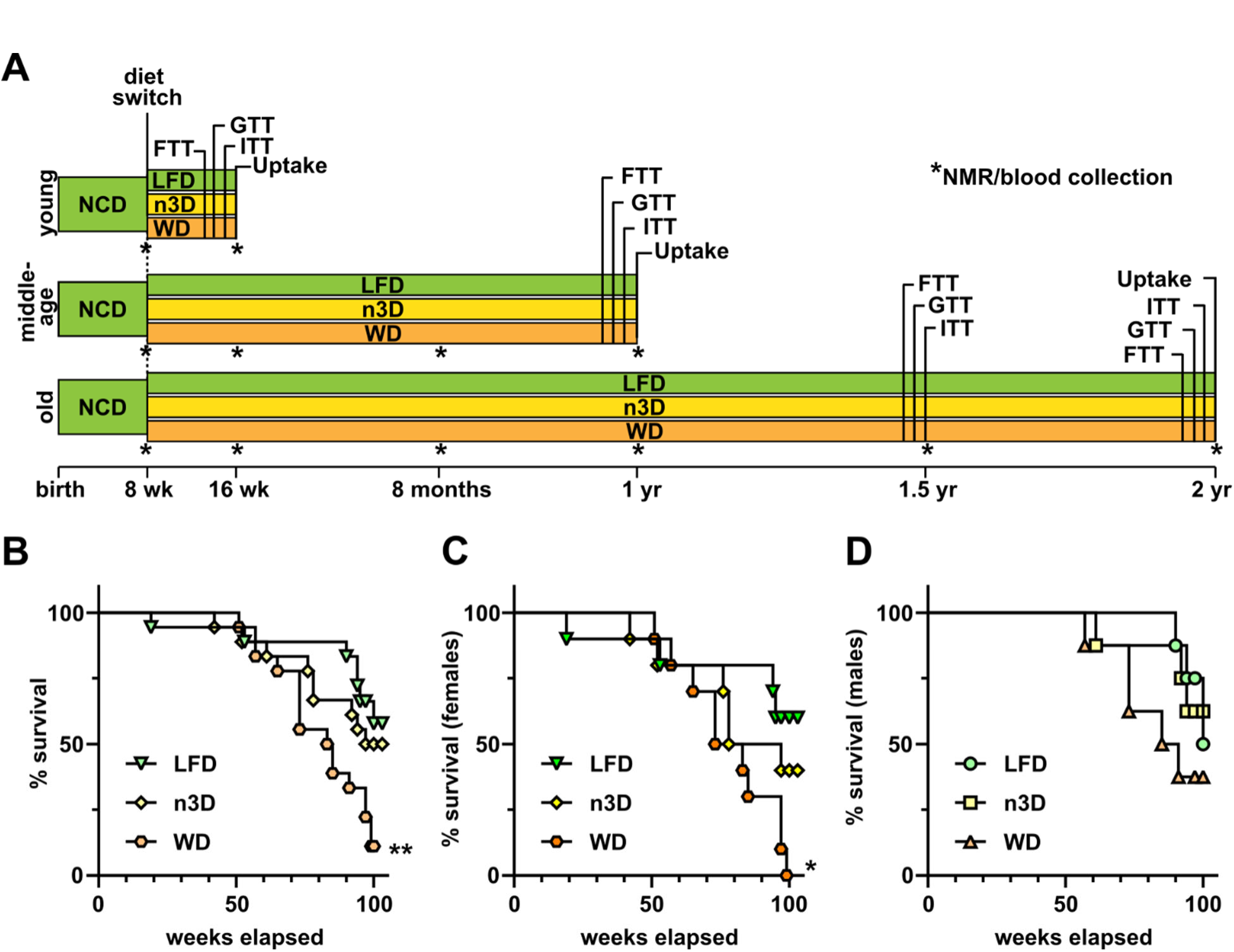
Experimental timeline and survival analysis. **A)** Timeline of experimental procedures for each age group. Each age group started with 8 female and 8 male mice per diet (the female 2-year cohorts started with 10 mice/cohort). Each cohort underwent fat tolerance tests (FTT) 3 weeks prior to sacrifice, glucose tolerance tests (GTTs) 2 weeks prior to sacrifice, and insulin tolerance tests (ITTs) 1 week prior to sacrifice. Uptake assays using radiolabeled triglyceride were performed as the terminal experiment. Fasting blood was collected and body mass analysis (NMR) performed at the indicated time points (*). **B-D)** Survival of combined female and male (B), female alone (C), and male alone (D) mice, in the 2-year cohort. Survival was analyzed using a log-rank (Mantel-Cox) test. *p<0.05, **p<0.01 compared to LFD mice. Complete lists of p values are listed in Supplementary Table 1.

### Effects of diets on lifespan in mice

In the 2-year cohort, a significant number of mice did not survive to the two-year mark. When we plotted survival, there was no difference in survival between LFD- and n3D-fed mice (**Fig. 1B**). However, survival was significantly reduced in mice fed a Western diet compared to those fed a LFD (**Fig. 1B**). When we separated mice by sex, survival was significant decreased in females fed a WD compared to LFD (**Fig. 1C**). Differences in male mice did not reach statistical significance (**Fig 1D**). Because of the low numbers of mice that survived to the 2-year mark on western diet, we decided to exclude the data from this time point in the main text. We have included data from the surviving mice in the 2-year cohort in the supplementary materials.

### Effects of diets on bodyweight, food consumption, and muscle composition in mice

After initiation of high fat feeding, we measured body weight weekly. As expected, males and females on the Western diet gained more weight than LFD fed mice (**Fig. 2**). Body weights of LFD- and WD-fed female mice increased through 1.5 years (**Fig. 2A**). In contrast, body weights of male mice fed either LFD or WD peaked at approximately 1 year of age and began to decline between ages 1 and 1.5 years (**Fig. 2B**). Not surprisingly, weights of WD-fed females dropped quickly as these mice began to die. Mice fed a diet high in omega-3 fatty acids had intermediate body weights. In males, n3D-fed mice had similar body weights to LFD-fed mice for the first 5 months on diet. Then body weights diverged, with n3D-fed mice gaining more weight until, by about a year and a half, body weights were similar to WD-fed mice. In female mice, body weights of n3D-fed mice diverged more quickly from LFD, but never approached the body weights of WD-fed mice. The lower weights of n3D-fed mice compared to WD-fed mice was likely due in part to lower food consumption. In both female and male mice, caloric intake of mice fed a n3D diet was not significantly different than those fed a LFD, whereas caloric intake was higher in mice fed a WD (**Fig. 2C, D**).

**Figure 2:**
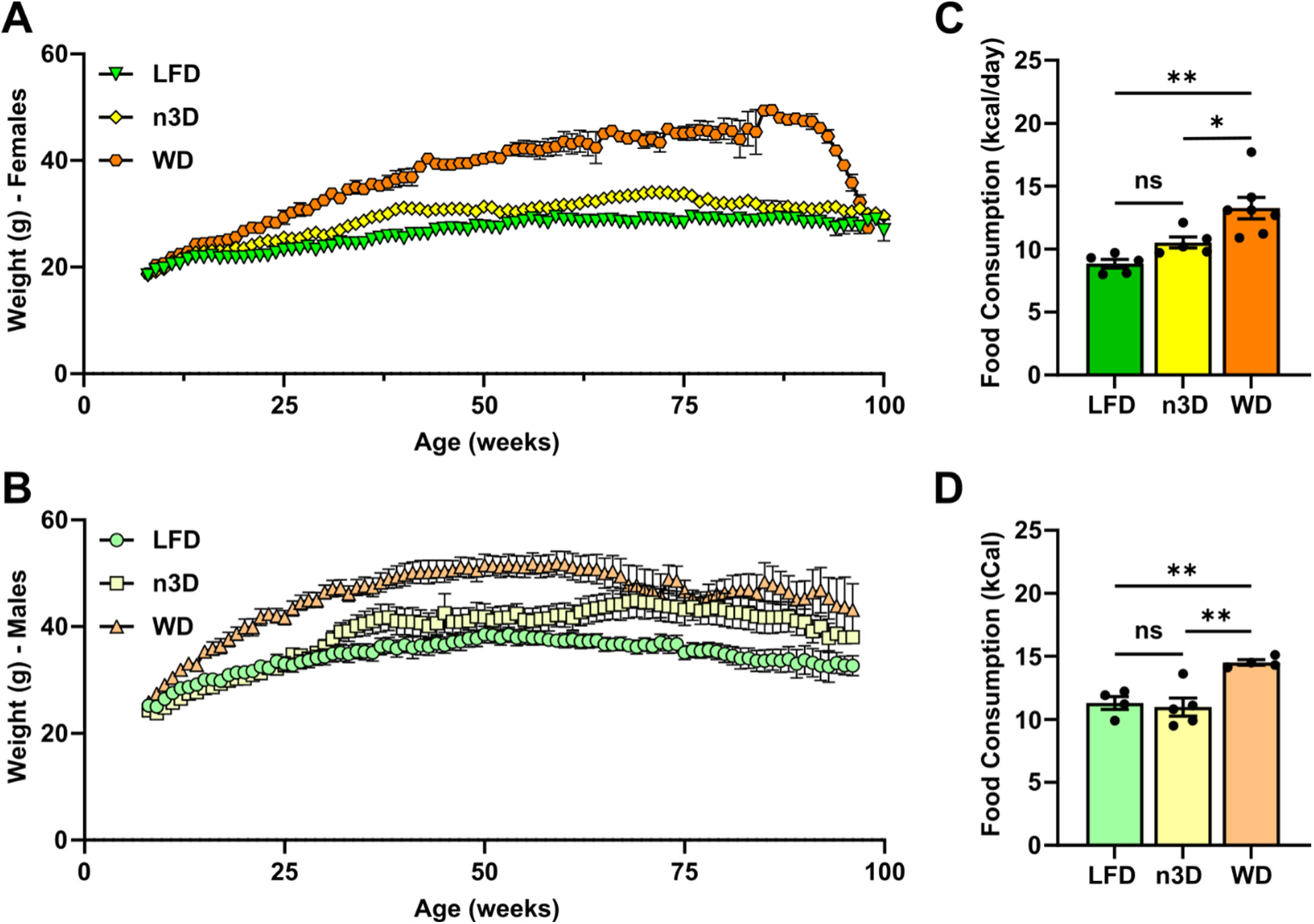
Body weight and food consumption in female and male mice. **A-B)** Body weights of female (A) and male (B) mice in the 2-year cohort, measured weekly. **C-D)** Food consumption of female (C) and male (D) mice. Food consumption was measured daily for 1 week and averaged. Mice were 22-47 weeks of age when food consumption was measured. ns=not significant, *p<0.05, **p<0.01 by Tukey’s multiple comparison test after 1-way ANOVA. Complete lists of p values are listed in Supplementary Table 1.

In both male and female mice fed a WD, the increased body weight was reflected in both a small increase in lean mass and a large increase in fat mass as measured by NMR (**Fig. 3, Fig. S1**). Surprisingly, in n3D-fed mice, the increased body weight compared to LFD-fed mice appeared to be due mostly to increased lean muscle mass. In females, fat mass of n3D-fed mice was never significantly higher than that in LFD-fed mice (**Fig. 3C**). In male mice, the mean fat mass of n3D-fed mice was slightly higher than in LFD, but this was only statistically significant at 8 months of age (**Fig. 3D**).

**Figure 3:**
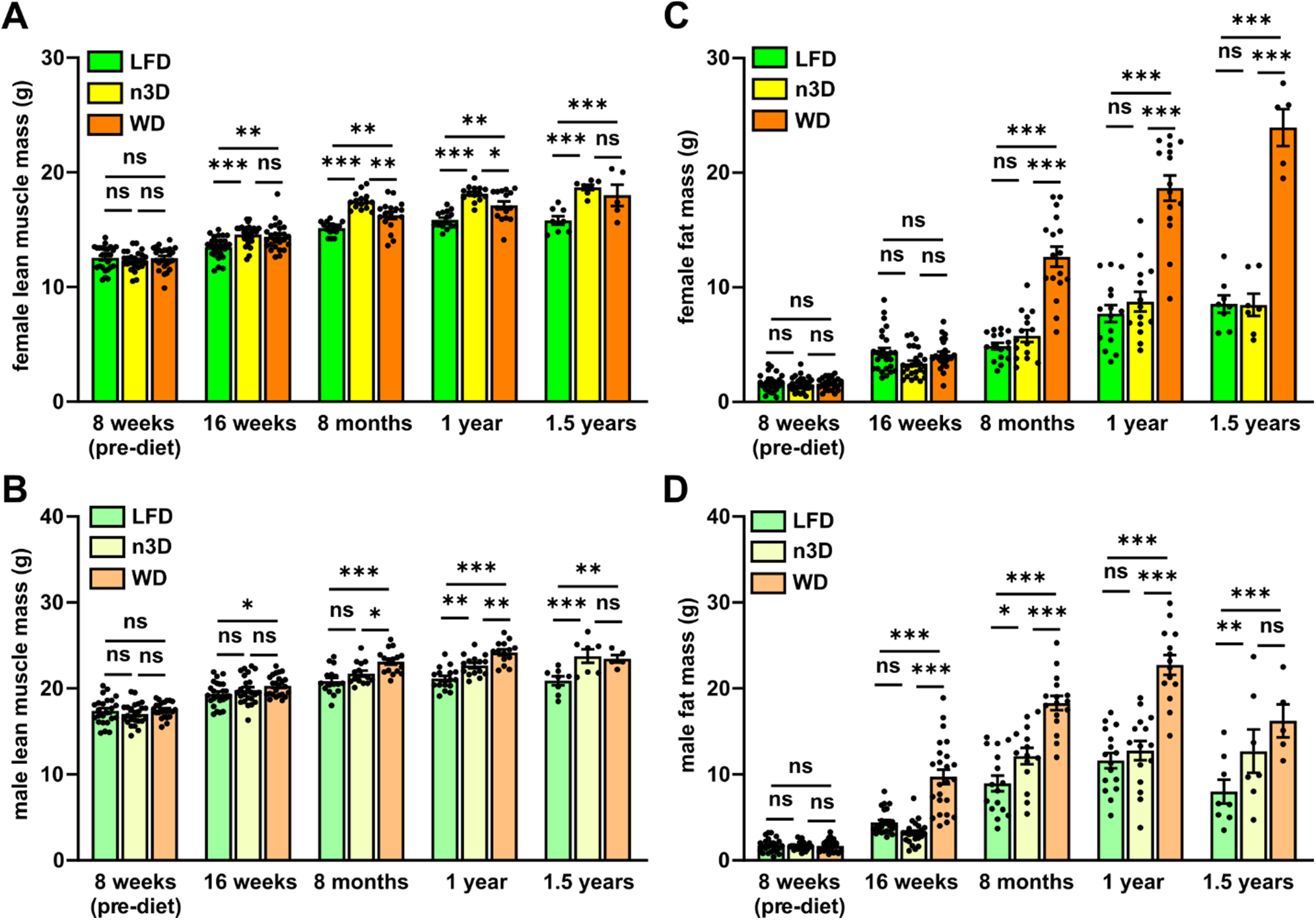
Body composition across age and diet. Lean muscle (**A-B**) and fat (**C-D**) mass in female (A,C) and male (B,D) mice as measured by NMR. ns=not significant, *p<0.05, **p<0.01, ***p<0.001 by Tukey’s multiple comparison test after 2-way ANOVA. Complete lists of p values are listed in Supplementary Table 1.

### Age- and diet-induced changes in triglyceride levels

In humans, fasting triglyceride levels increase with age.^19–21^ To determine if plasma triglycerides in mice also change with age, we measured triglycerides in our two-year cohort at 8, 16, 26, 52, 78, and 100 weeks. Contrary to our expectations, fasting plasma triglyceride levels in females on all diets decreased until 1.5 years of age (**Fig. 4A**). For mice that survived to 100 weeks, plasma triglycerides levels at 100 weeks were similar to those at 78 weeks (**Fig. S2A**). Plasma triglyceride levels in female LFD and WD fed mice were not different, but n3D-fed mice had modestly, but significantly lower TGs (p=0.001 by 2-way ANOVA)(**Fig. 4A**). Regardless of diet, in male mice fasting plasma triglyceride levels decreased until 26 weeks of age and then remained steady until 78 weeks (**Fig. 4B**). Male mice on n3D that survived to 2 years showed a rise in TG levels between 1.5 and 2 years (**Fig. S2B**).

**Figure 4:**
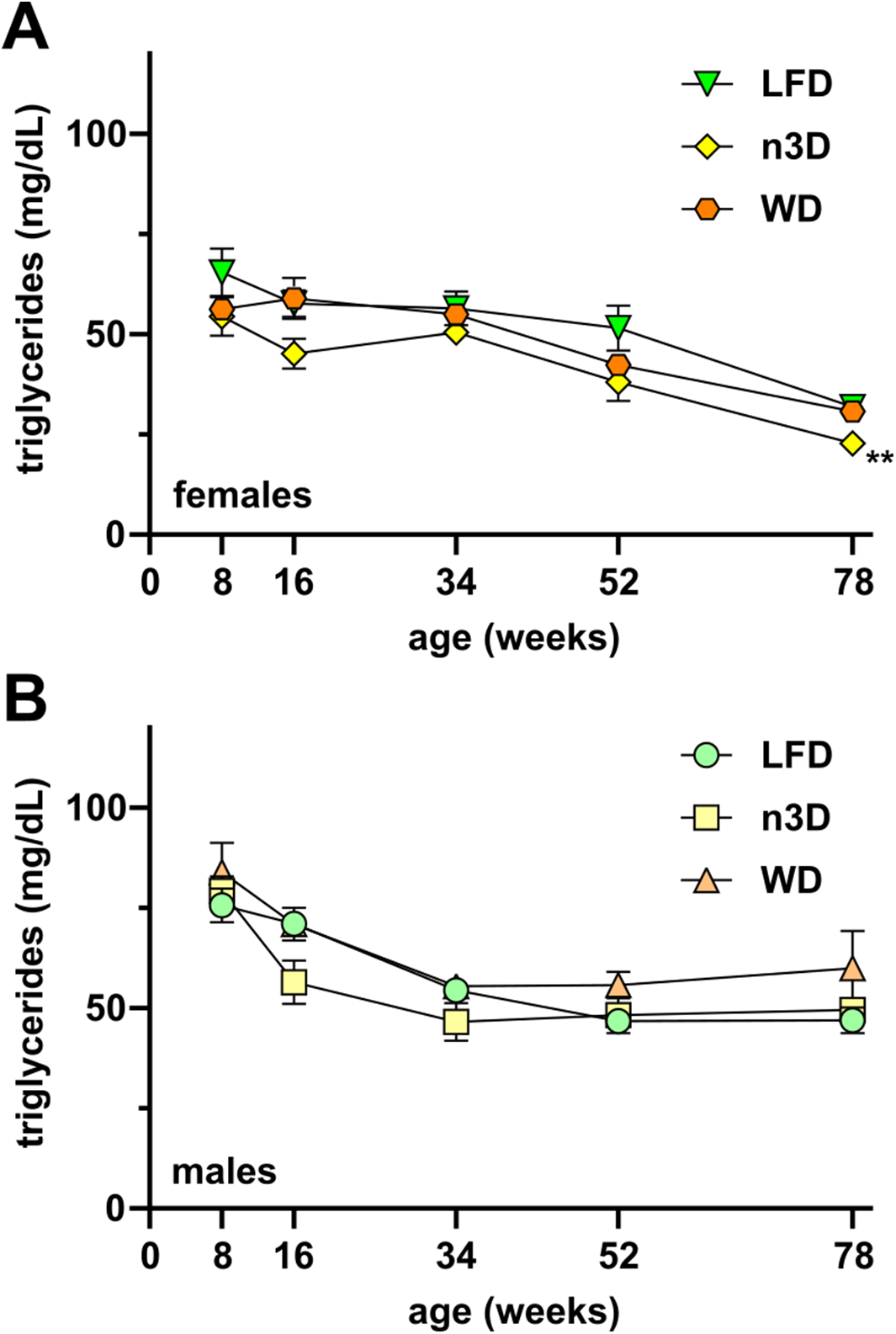
Fasting plasma triglyceride levels across age and diet. Plasma was collected from mice following a 6 h fast and plasma triglyceride levels were measured in female **(A)** and male **(B)** mice. **p<0.01 by mixed effects analysis. Complete lists of p values are listed in Supplementary Table 1.

### Age- and diet-induced changes in oral fat tolerance

Previous studies with aged human subjects found that clearance of serum lipid levels after a fat-loaded meal was strikingly delayed in older subjects compared with younger subjects.^19,22,23^ To elucidate if a similar delay occurs in aged mice, we performed oral fat tolerance tests (FTTs) in all 3 diet groups at 13 weeks (5 weeks on diet), 49 weeks, 75 weeks, and 2 years. In 13-week-old mice, we observed the expected peak in triglyceride levels 2-3 hours after oral gavage and subsequent return of triglyceride levels towards baseline (Fig. 5A,B). In these younger mice, both the n3D and WD-fed male and female mice had increased triglyceride levels compared to LFD-fed mice, but this difference was not statistically significant. Surprisingly, and contrary to our hypothesis, the large spike in plasma TG levels following gavage was largely absent in all of the older groups (**Fig. 5C-F; Fig. S3**). Interestingly, in the female mice of the older cohorts, TG levels after oral gavage were significantly higher in n3D-fed mice than those of LFD or WD fed mice (**Fig. 5C,E**).

**Figure 5:**
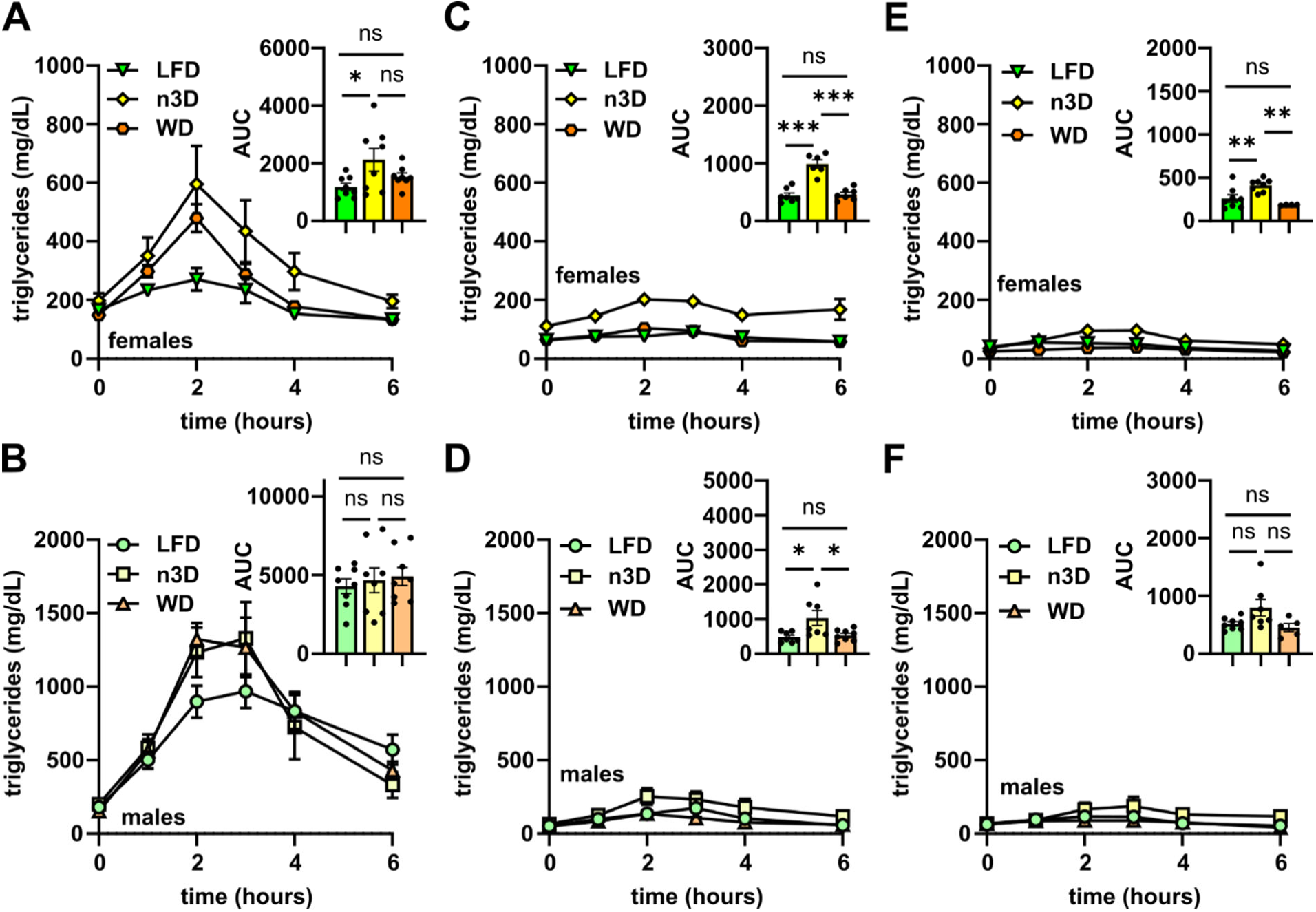
Oral fat tolerance tests. Mice were fasted (12 h) and plasma was collected before and 1,2,3,4 and 6 h after olive oil gavage (10 µl/g body weight). Plasma triglyceride levels were measured at each time point in 13 week old female **(A)** and male **(B)** mice, 49-week old female **(C)** and male **(D)** mice, and 75-week old female **(E)** male **(F)** mice. Insets show area under the curve (AUC). ns=not significant, *p<0.05, **p<0.01, ***p<0.001 by Tukey’s multiple comparison test after 1-way ANOVA. Complete lists of p values are listed in Supplementary Table 1.

### Effects of diets and age on triglyceride clearance and tissue triglyceride uptake

The partitioning of triglycerides between different adipose tissue depots and other peripheral locations differs not only between sexes, but also during aging.^27–32^ To better understand how sex, age, and diet can lead to alterations in triglyceride storage and partitioning we performed plasma triglyceride clearance and tissue uptake assays. Mice were injected with chylomicrons containing radiolabeled triglyceride, and the clearance of radiolabel from the bloodstream was measured over 15 minutes. After 15 minutes, mice were treated with Tyloxapol to block further lipolysis of lipoproteins and tissues were collected and assessed for radiolabel uptake. The profile of triglyceride clearance from the blood was similar across all ages (**Fig. 6; Fig S4**). While there were no diet-induced differences in clearance in female mice (**Fig. 6A,C**) nor in 16 week old male mice (**Fig. 6B**), in 1 year-old male mice, WD-fed mice cleared triglycerides faster than LFD- or n3D-fed mice (**Fig. 6D**).

**Figure 6:**
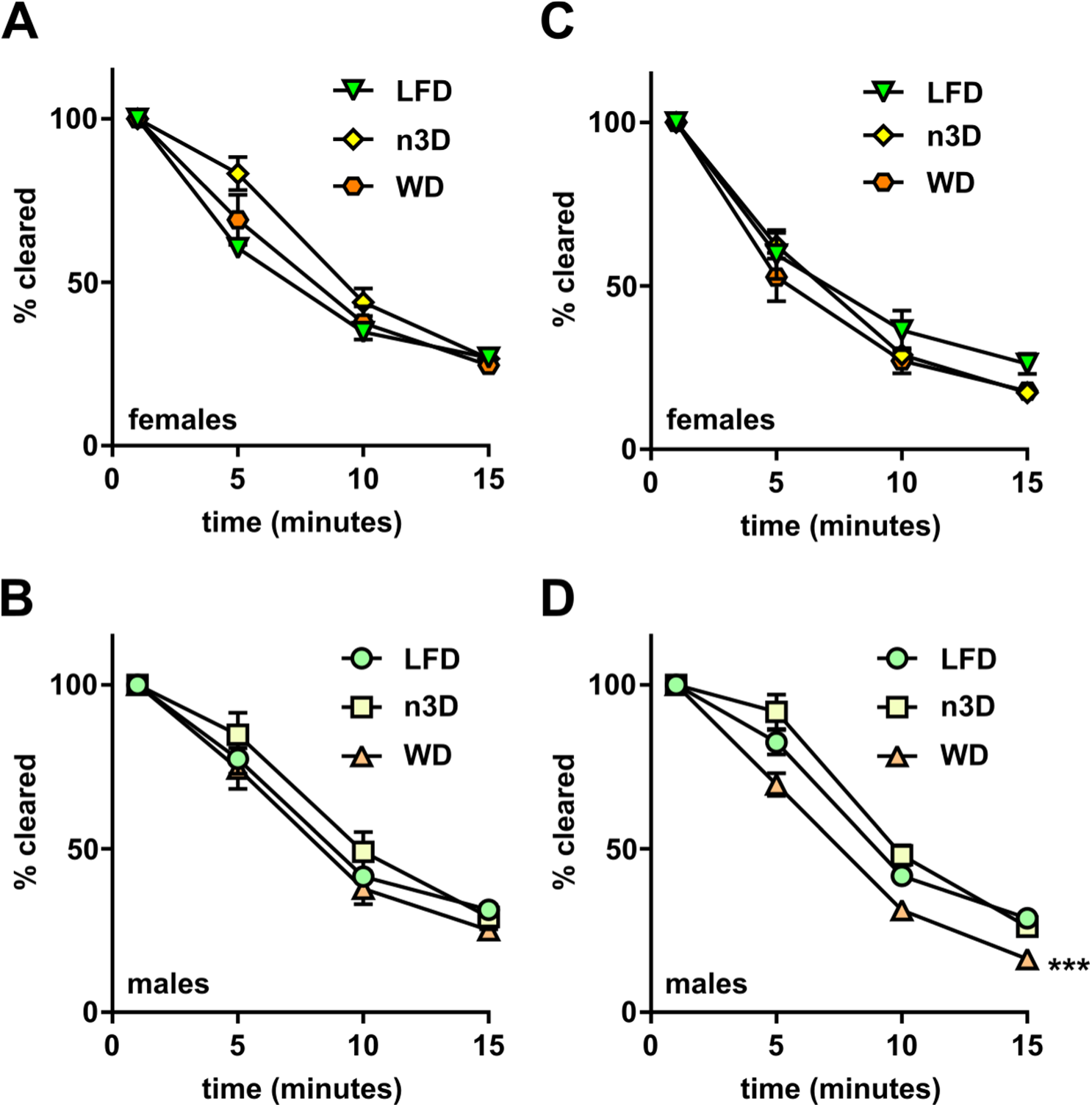
Plasma triglyceride clearance. Mice were fasted (6 h) and injected intravenously with ^3^H-triglyceride containing chylomicrons. Blood was collected after 1, 5, 10, and 15 min after injection to measure clearance of radiolabel from the plasma. Points represent radiolabel remaining in the plasma as a percentage of the 1-min time point (means ± SEM) in 16-week-old female **(A)**, 16-week-old male **(B)**, 1-year-old female **(C)**, and 1 year old male **(D)** mice. ***p<0.001 for diet by repeated measures 2-way ANOVA. Complete lists of p values are listed in Supplementary Table 1.

With the exception of the quadriceps in LFD-fed females, on a per gram tissue basis, triglyceride uptake either stayed the same or decreased from 16 weeks to 1 year in all tissues and on all diets (**Fig. 7,8**). The most significant age-mediated decreases in TG uptake were seen in white adipose tissue, particularly those fed high-fat diets (**Fig. 7E,F**; **Fig 8E,F**). It is important to note that even though uptake went down on a per gram basis in many tissues, because of the increased size of these tissues with age, the total amount of TG entering these tissues either stayed the same or increased (**Fig. S5**). The most striking effects of diet on triglyceride uptake were found in the heart and in the liver. In both male and female hearts, n3D-fed mice took up substantially more radiolabel than either LFD- or WD-fed mice (**Fig. 7A, 8A**). This was true across age groups and was true whether calculating uptake on a per gram of tissue (**Fig. 7A, 8A**) or a total tissue (**Fig. S5**) basis. In the liver, reduced TG uptake (on a per gram tissue basis) was observed for both high fat diets compared to the LFD (**Fig. 7B, 8B**), however these differences were less striking when calculating uptake on a whole tissue basis (**Fig. S5**). When analyzing the surviving 2-year-old mice, TG uptake did go up in some tissues, but again the low numbers, questionable health status, and lack of surviving WD-fed female mice in these cohorts make it difficult to interpret the data (**Fig. S6, S7**).

**Figure 7:**
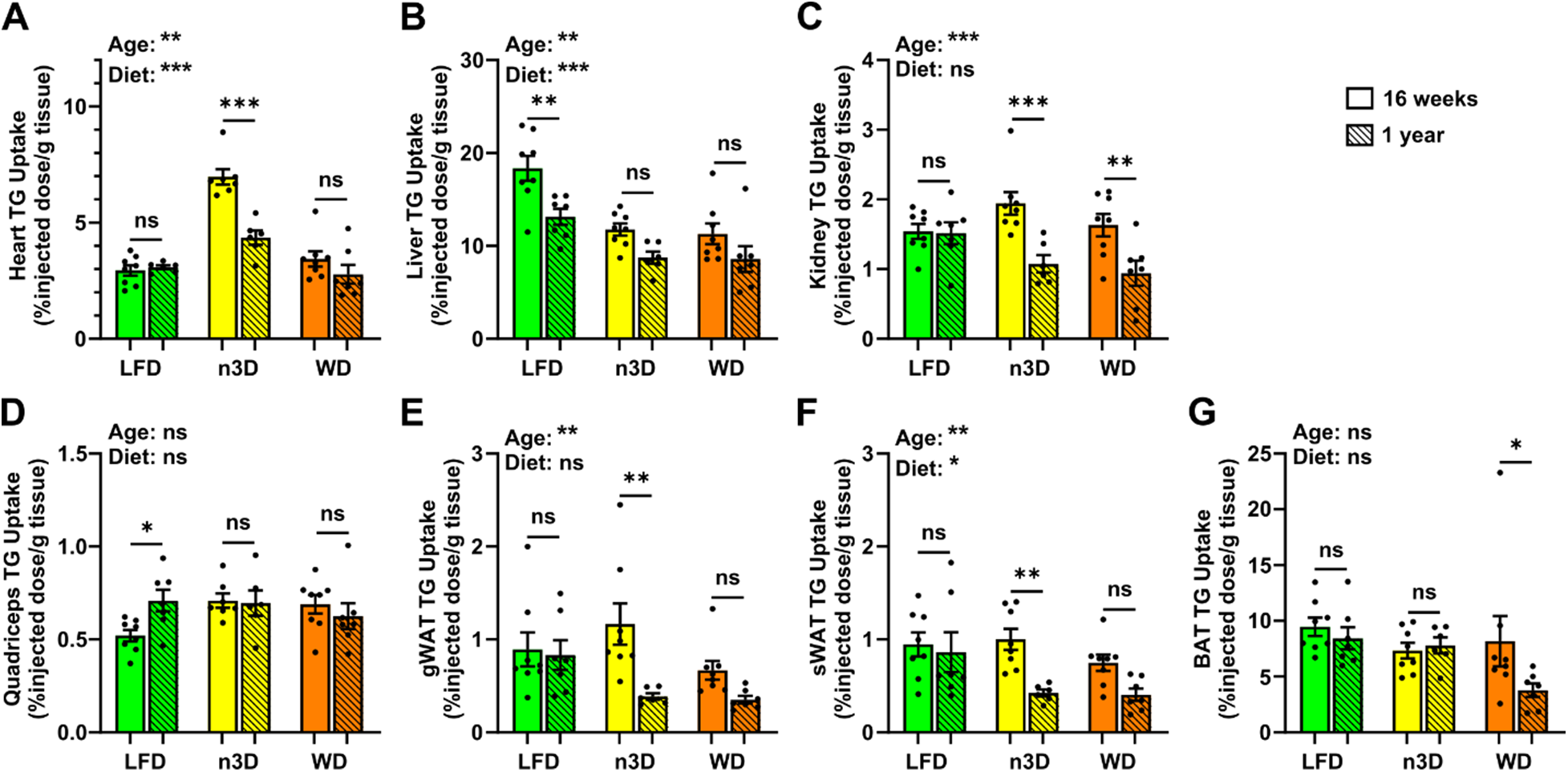
Tissue triglyceride uptake in female mice. At 16 weeks or 1 year of age female mice were fasted (6 h) and injected intravenously with ^3^H-triglyceride containing chylomicrons. After 15 minutes, tissues were harvested and uptake of radiolabel (% injected dose/g tissue) was measured in heart **(A)**, liver **(B)**, kidney **(C)**, quadriceps muscle **(D)**, gonadal white adipose tissue (gWAT)**(E)**, subcutaneous adipose tissue (sWAT)**(F)**, and brown adipose tissue (BAT)**(G)**. Bars represent mean ± SEM. ns=not significant, *p<0.05, **p<0.01, ***p<0.001 by 2-way ANOVA and subsequent Tukey’s multiple comparison test. Complete lists of p values are listed in Supplementary Table 1.

**Figure 8:**
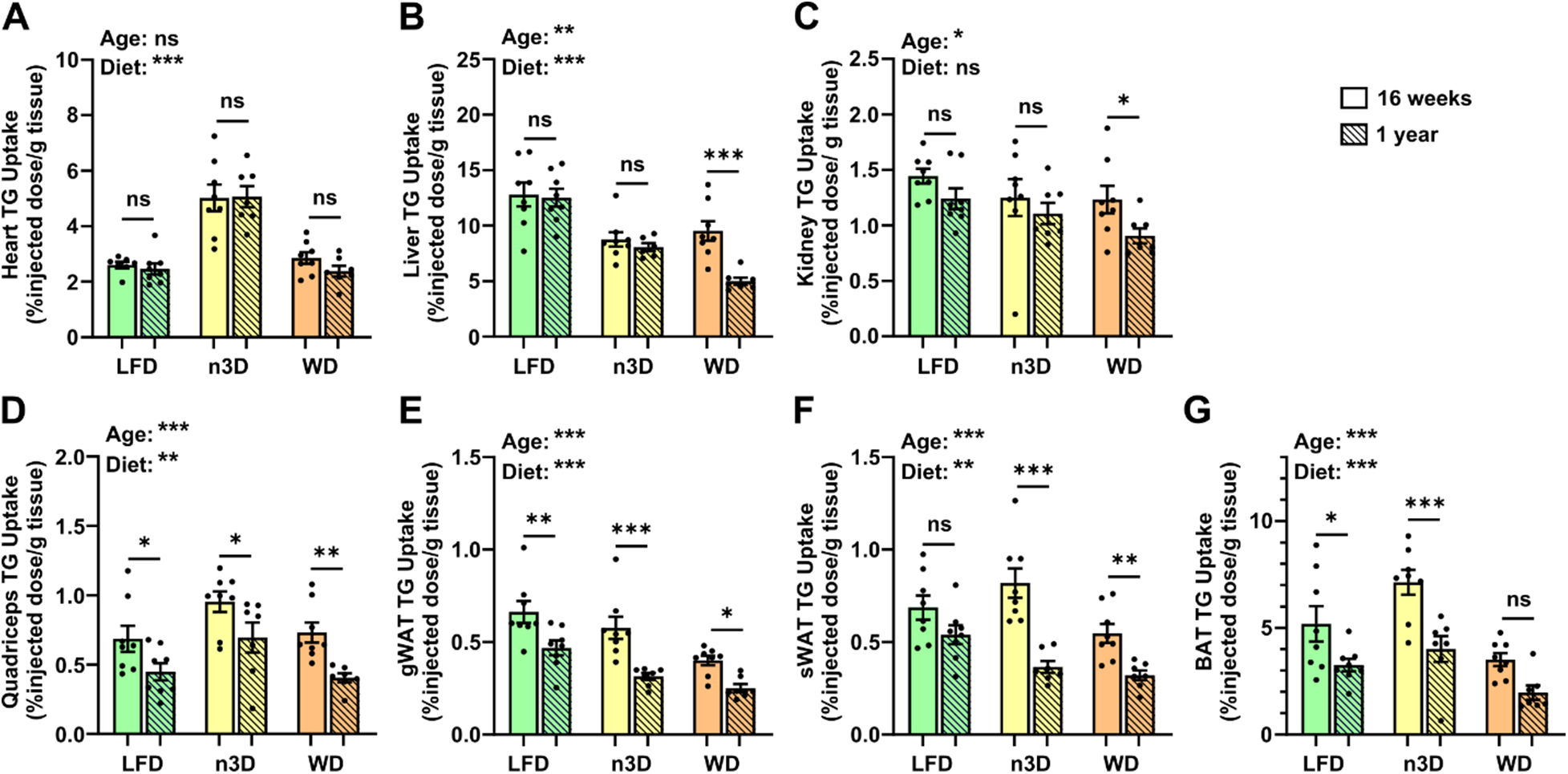
Tissue triglyceride uptake in male mice. At 16 weeks or 1 year of age mice were fasted (6 h) and injected intravenously with ^3^H-triglyceride containing chylomicrons. After 15 minutes, tissues were harvested and uptake of radiolabel (% injected dose/g tissue) was measured in heart **(A)**, liver **(B)**, kidney **(C)**, quadriceps muscle **(D)**, gonadal white adipose tissue (gWAT)**(E)**, subcutaneous adipose tissue (sWAT)**(F)**, and brown adipose tissue (BAT)**(G)**. Bars represent mean ± SEM. ns=not significant, *p<0.05, **p<0.01, ***p<0.001 by 2-way ANOVA and subsequent Tukey’s multiple comparison test. Complete lists of p values are listed in Supplementary Table 1.

### Effects of diets and age on glucose metabolism

To understand how diet and age effect glucose homeostasis in mice, we performed glucose tolerance tests (GTTs) and insulin tolerance tests (ITTs) at 14-15 weeks (6-7 weeks on diet), 50-51 weeks, 76-77 weeks, and 2 years of age in both males and females. After 6 weeks on WD, there was a significant loss of glucose tolerance compared to LFD in both male and female mice (**Fig. 9A,D**). Female n3D-fed mice had reduced glucose tolerance, but male n3D-fed mice were not significantly different from LFD-fed mice (**Fig. 9A**). Surprisingly, after 1 year, there were no statistical differences in glucose tolerance across diets (**Fig. 9B,E**), though it should be noted that by 1.5 year there was substantial variability in the WD-fed group (**Fig. 9C,F**).

**Figure 9:**
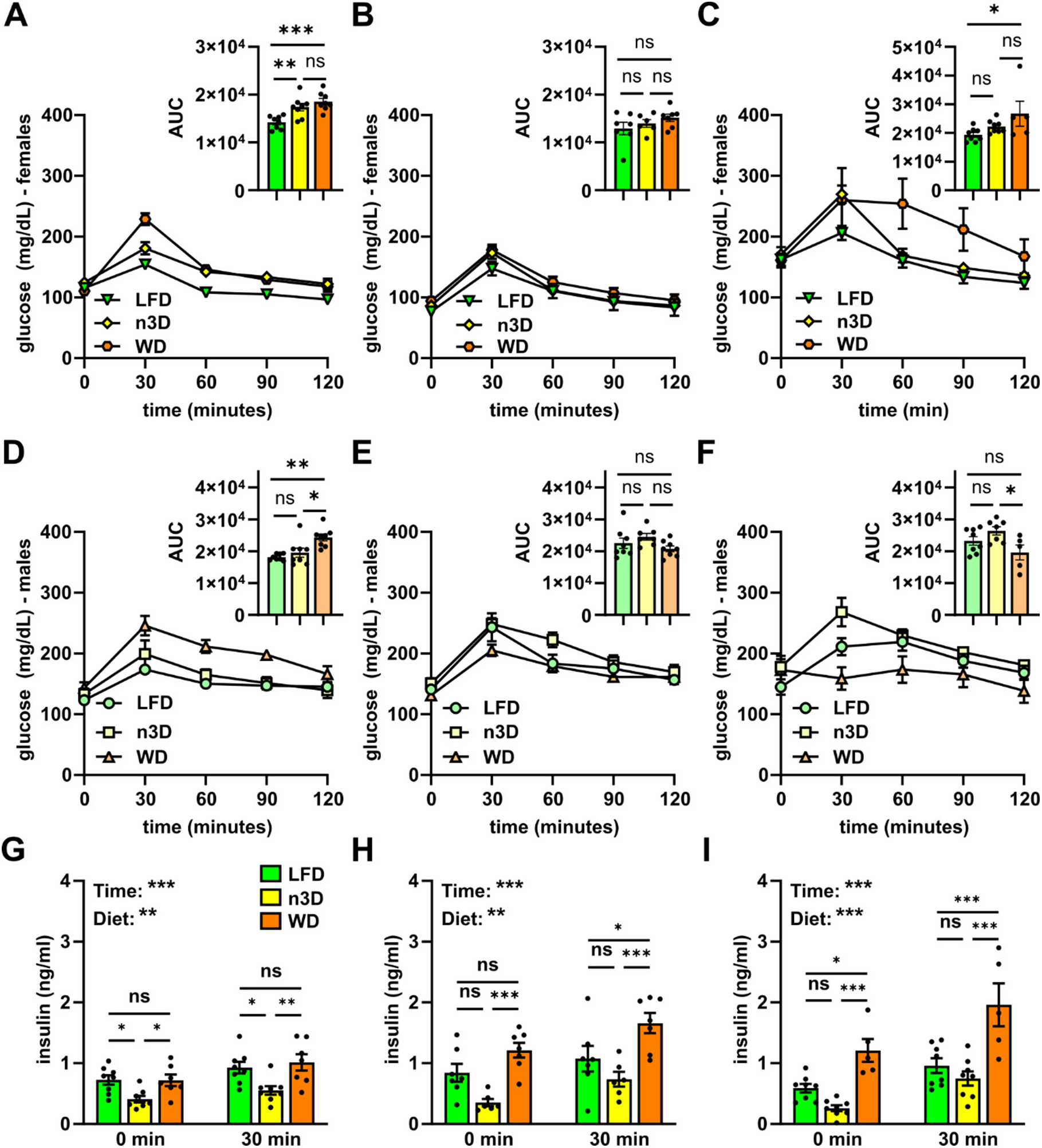
Glucose tolerance tests and plasma insulin levels. **A-F)** Glucose tolerance tests were performed on fasted (6 h) female (A-C) and male (D-F) mice at 14 (A,D), 50 (B,E), and 76 (C,F) weeks of age. Mice were injected with glucose (1 g/kg) and blood glucose concentrations were measured over 2 h. Points represent glucose levels (means ± SEM) at each respective time point. Insets show area under the curve (AUC). ns=not significant, *p<0.05, **p<0.01, ***p<0.001 by Tukey’s multiple comparison test after 1-way ANOVA. **G-I)** Plasma insulin levels before and 30 minutes after glucose injection in 14-(G), 50-(H), and 76- (I) week-old female mice. Bars represent mean ± SEM. ns=not significant, *p<0.05, **p<0.01, ***p<0.001 by 2-way ANOVA and subsequent Tukey’s multiple comparison test. Complete lists of p values are listed in Supplementary Table 1.

As part of our GTT assays, we measured insulin levels in female mice before and 30 minutes after glucose injection. Interestingly, n3D-fed mice had lower insulin levels both prior to and 30 minutes after glucose injection, especially compared to WD fed mice (**Fig 9G-I**). This was true at all ages (**Fig 9G-I**, **Fig S8**), though mouse numbers were low at 2 years. The reduced insulin levels in n3D-fed mice, despite similar glucose tolerance, suggested that these mice might have increased insulin tolerance. Indeed, insulin tolerance tests suggested that n3D-fed mice were more insulin sensitive than mice fed the other two diets (Fig. 10; Fig S9). Increased insulin sensitivity in n3D-fed mice was also supported by the fact that when performing ITTs on 1.5-year-old female mice, only 2 of the 8 n3D-fed mice were able to finish the time course as all others had to be given a bolus of glucose due to hypoglycemia. 3 of 8 LFD-fed mice also had to be given glucose.

**Figure 10:**
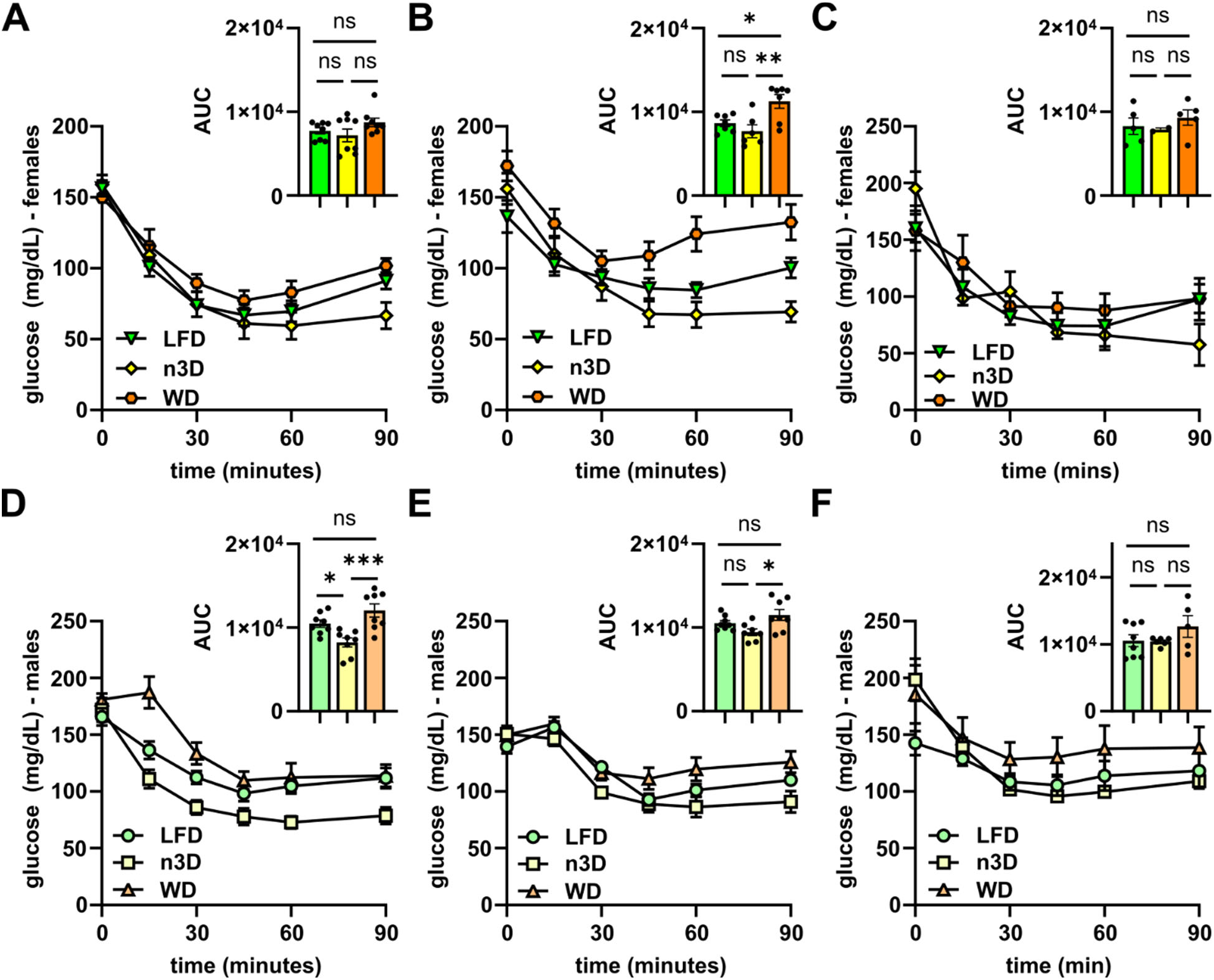
Insulin tolerance tests. Insulin tolerance tests were performed in fasted (4 h) female **(A-C)** and male **(D-F)** mice at 15 (A,D), 51 (B,E), or 77 (C, F) weeks of age. Mice were injected with 0.5 U/mL of human insulin (Humalin-R) and blood glucose concentrations were measured over 90 min. Points represent glucose levels (means ± SEM) at each respective time point. Insets show area under the curve (AUC). ns=not significant, *p<0.05, **p<0.01, ***p<0.001 by Tukey’s multiple comparison test after 1-way ANOVA. Complete lists of p values are listed in Supplementary Table 1.

## Discussion

In this study we fed mice three different diets for up to two years to examine how age and diet affected triglyceride metabolism in mice. Our major findings include 1) a high-fat western diet reduces life span; 2) whereas in humans plasma triglyceride levels increase with age and triglyceride clearance after a fatty meal decreases, the opposite appears to be true in mice where both fasting triglyceride levels and postprandial spikes in plasma TG levels were lower in older mice than those in young mice; and 3) a diet high in polyunsaturated and omega-3 fat increased the uptake of triglyceride-derived fatty acid into the heart and led to an increase in lean muscle mass.

A western diet is known to exacerbate several metabolic diseases in humans and is thought to lower mortality.^33–35^ In mice, a western diet has been shown to increase mortality in combination with induced pathologies such as sepsis,^36^ hepatic steatosis,^37^ and necrotising pancreatitis.^38^ Here we show that western diet alone, in the absence of any other insult, decreased survival in mice when compared to a low-fat diet. Interestingly this change in survival was more pronounced in female mice than in male mice.

A major goal of this study was to understand how triglyceride metabolism changes with age in mice and compare these changes to those previously observed in humans. In humans, both fasting and postprandial TG levels are greater in men than in women, and in both sexes TG levels increase with age.^20^ Although we did find that fasting and postprandial TG levels were higher in male mice than female mice, mostly notably at younger ages, unlike humans, plasma triglyceride levels did not increase with age. Instead, we found that for both male and female mice, fasting triglyceride levels were lower in old mice versus young mice. This was consistent with the findings of *Houtkooper et al* who found increased free fatty acids and decreased triglyceride levels in aged (22 months) mice compared to young (3 months) mice,^39^ but differed from the findings of Xiong et al. who found that plasma TG levels were greater in 20 month-old mice than 2-month-old mice.^40^ Interestingly, we also found that in our mice, TG levels decreased with age regardless of diet and mice fed a western diet did not have significantly higher TG levels than those fed a low-fat diet.

As with fasting TG levels, postprandial TG phenotypes also differed between our mouse cohort and what has been observed in humans. Several human studies have shown that the post-prandial increase in TG levels after a high fat meal is both higher and longer in aged individuals than in young individuals.^19,22,23^ Here we found that older male and female mice had a much lower rise in plasma TG after a high fat gavage. This lower TG rise could be mediated by lower fat absorption and subsequent secretion of chylomicrons into the circulation, or it could be mediated by faster triglyceride clearance. Yamamoto et al. also found that the postprandial increase in plasma triglycerides was lower in aged mice and concluded that absorption was lower based on finding lower levels of pancreatic lipase activity.^41^ It should be noted that in that study, plasma TG clearance was not measured, postprandial TG levels were only measured at the 3 h mark, and experiments were only done in male mice.^41^ Despite the limitations of this previous study, our data also suggest a decrease in fat absorption in both male and female aged mice, as our triglyceride clearance assays did not show a significant difference in TG clearance rate with age (see Fig. 6). Interestingly, humans also seem to have decreased fat absorption rates with age,^42,43^ suggesting that the increase in postprandial TG levels in humans is likely due to dysfunctional clearance. Indeed, Vinagre et al. found that LPL activity in post-heparin plasma was reduced in aged individuals and that after injection of chylomicron-like emulsions, clearance of remnant particles was reduced.^23^ Overall, our data suggest that mice may have a similar decrease in fat absorption when compared to humans, but that mice maintain functional TG clearance with age whereas humans do not. As a result, post-prandial triglyceride levels increase with age in humans, but decrease with age in mice.

Some of the most intriguing findings of this study come from the cohort of mice fed a diet high in polyunsaturated and omega-3 fatty acids. Chronic low-grade inflammation is associated with aging^44^ and obesity.^45^ This heightened inflammatory state presents a potential therapeutic target for disturbances in glucose and lipid metabolism during aging. Omega-3 polyunsaturated fatty acids have anti-inflammatory properties and might prove to be a therapeutic agent.^46^ A meta-analysis found omega-3 treatment reduced C-reactive protein and IL-6 plasma levels in adults 45 or older.^47^ Low consumption of seafood-derived omega-3 fats is associated with increased risk of diet-related cardiometabolic death,^48^ while higher circulating levels of omega-3 fatty acids are associated with lower total mortality and lower risk in cardiovascular deaths in older adults.^49^ Omega-3 fats have becoming increasingly studied for their role in metabolic disease; however, few studies have examined long term treatment with Omega-3 fatty acids and metabolic health in mice. We found that mice fed the n-3 fat diet had lower body weights than mice fed a western diet, and unlike WD-fed mice did not have significantly reduced survival compared to LFD-fed mice. Mice fed an n-3 fatty acid diet, particularly female mice, had increased lean mass, but not increased fat mass compared to LFD-fed mice. These mice also appeared to be more insulin sensitive and have lower insulin levels than LFD- or WD-fed mice. Moreover, both male and female mice fed the n-3 fatty acid diet had a striking increase in triglyceride uptake into the heart. The low body weights in n-3 fat-fed mice are likely a result of the lower calorie-consumption observed in these mice compared to WD-fed mice. Likewise, the improved survival, lower insulin levels, and improved insulin sensitivity could be ascribed to the lower calorie consumption and the anti-inflammatory effects of omega-3 fatty acids. However, it is mechanistically unclear how an n-3 fatty acid diet leads to increased fatty acid uptake into the heart and an increase in lean mass rather than fat mass. It is also unclear if these two phenotypes are related to each other or to any of the other metabolic phenotypes observed in these mice. It is possible that the increased uptake into heart drove increased lean muscle mass. After 1 year on diet the hearts of female mice fed the n-3 diet were significantly larger than those is LFD fed mice (170.8 vs 137.1 mg; p=0.004). However, hearts in male mice and skeletal muscle in either sex were not significantly different.

Our study has several limitations. Although we performed assays on two-year old mice, the number of mice that died before reaching two years of age left the assays performed at this time point underpowered. Moreover, although we found that mice fed a western diet had reduced survival, we did not investigate the causes of death in these mice. Another limitation is that we examined only three diets. Although we carefully chose the diets used in this study, there are, of course, several different types of high-fat diets utilized in studies of diet-induced obesity and insulin resistance. It is possible that certain diets could have very different effects on triglyceride metabolism. Finally, there are a number of metabolic parameters that we did not measure. For example, although our data suggest that fat absorption decreases with age, we did not directly measure fat absorption. A more complete measurement of different metabolic parameters as well as gene and protein expression could provide a more comprehensive picture of age- and diet-induced changes in triglyceride metabolism.

## Acknowledgements

We thank Kelli Sylvers-Davie and Sydney Walker for technical assistance.

## Non-standard Abbreviations and Acronyms

FTT: fat tolerance test
GTT: glucose tolerance test
ITT: insulin tolerance test
LFD: low-fat diet
n-3: omega-3
n3D: omega-3 enriched diet
TG: triglyceride
WD: Western diet

