## Supplemental Data for "Effects of Age and Diet on Triglyceride Metabolism in Mice"

Short Title: Age, Diet, and Triglyceride Metabolism

**Supplementary Figure 1**

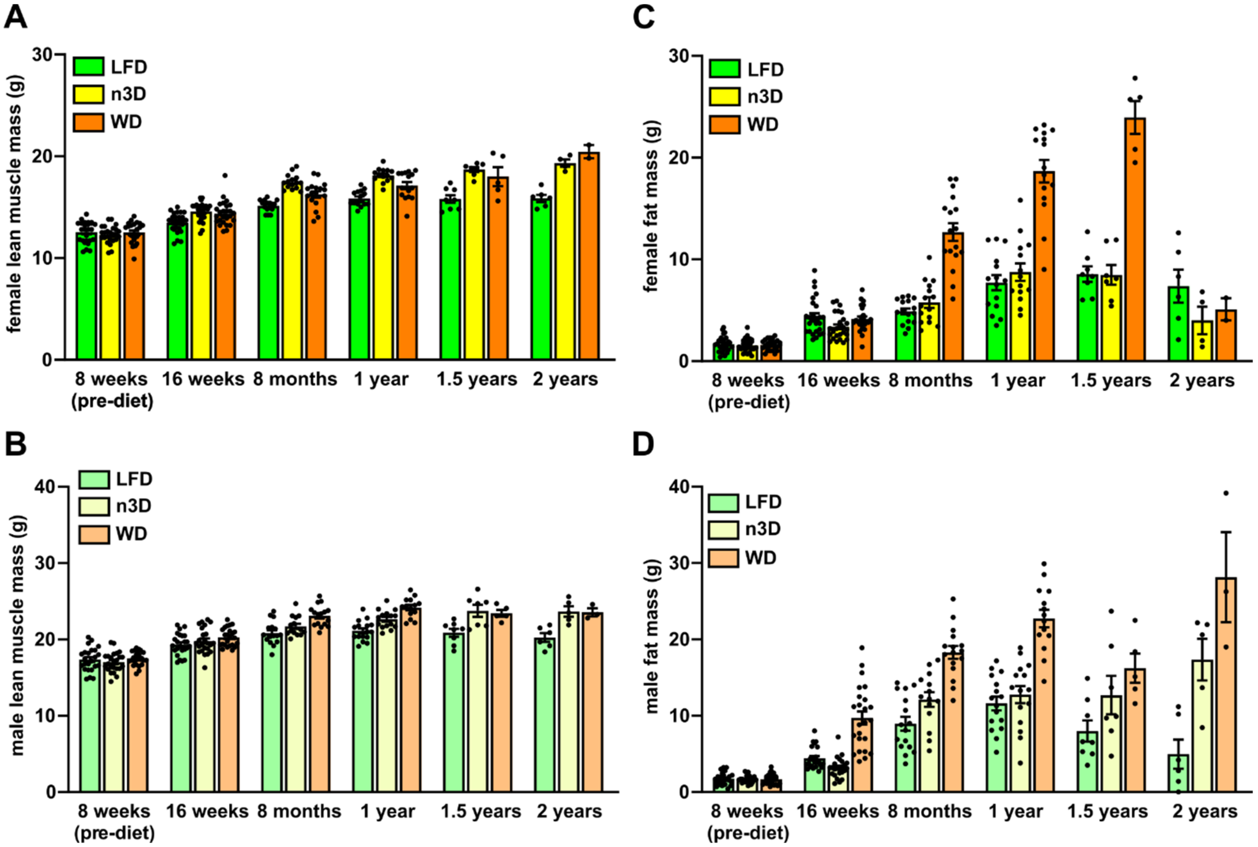


**Supplementary Figure 1: Body composition across age and diet.** Lean muscle (A-B) and fat (C-D) mass in female (A,C) and male (B,D) mice as measured by NMR.

**Supplementary Figure 2**

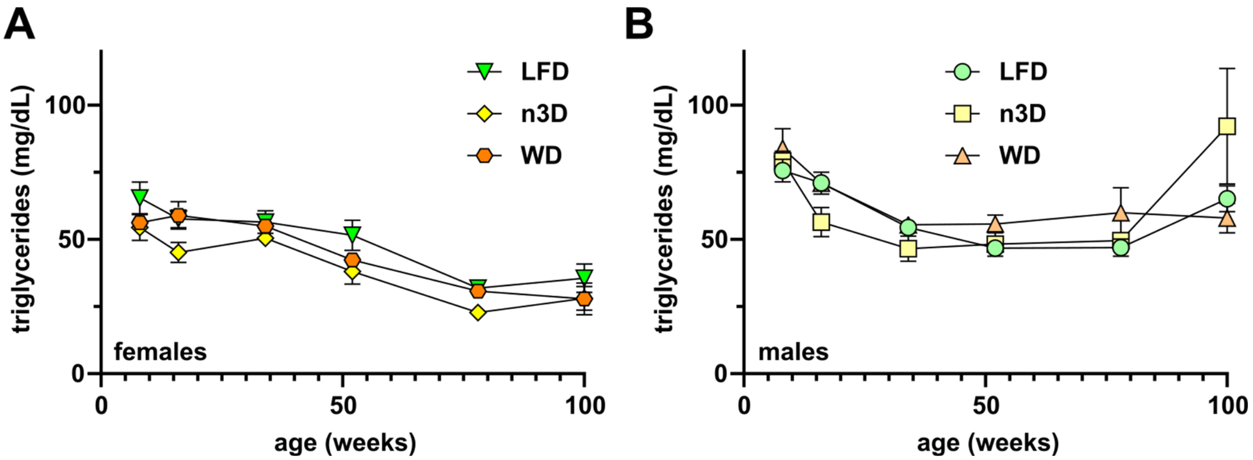


**Supplementary Figure 2: Fasting plasma triglyceride levels across age and diet.** Plasma was collected from mice following a 6 h fast and plasma triglyceride levels were measured in female **(A)** and male **(B)** mice.

**Supplementary Figure 3**

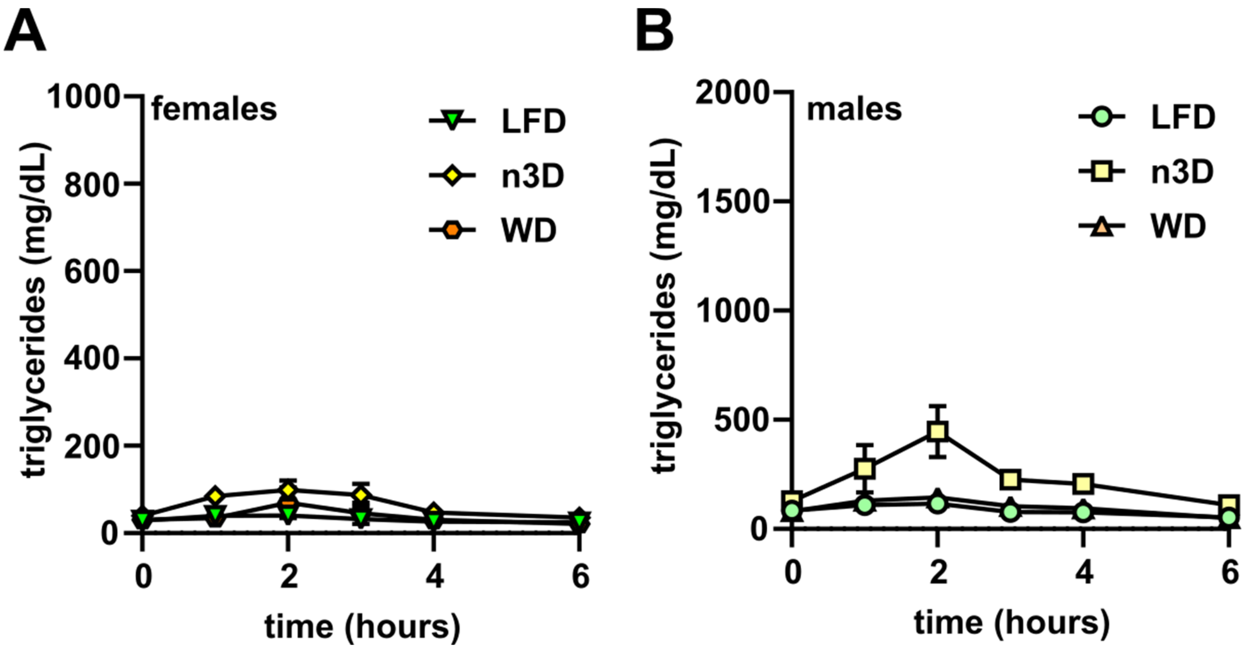


**Supplementary Figure 3: Oral fat tolerance tests in old mice.** Mice were fasted (12 h) and plasma was collected before and 1,2,3,4 and 6 h after olive oil gavage (10 µl/g body weight). Plasma triglyceride levels were measured at each time point in 2-year old female **(A)** and male **(B)** mice.

**Supplementary Figure 4**

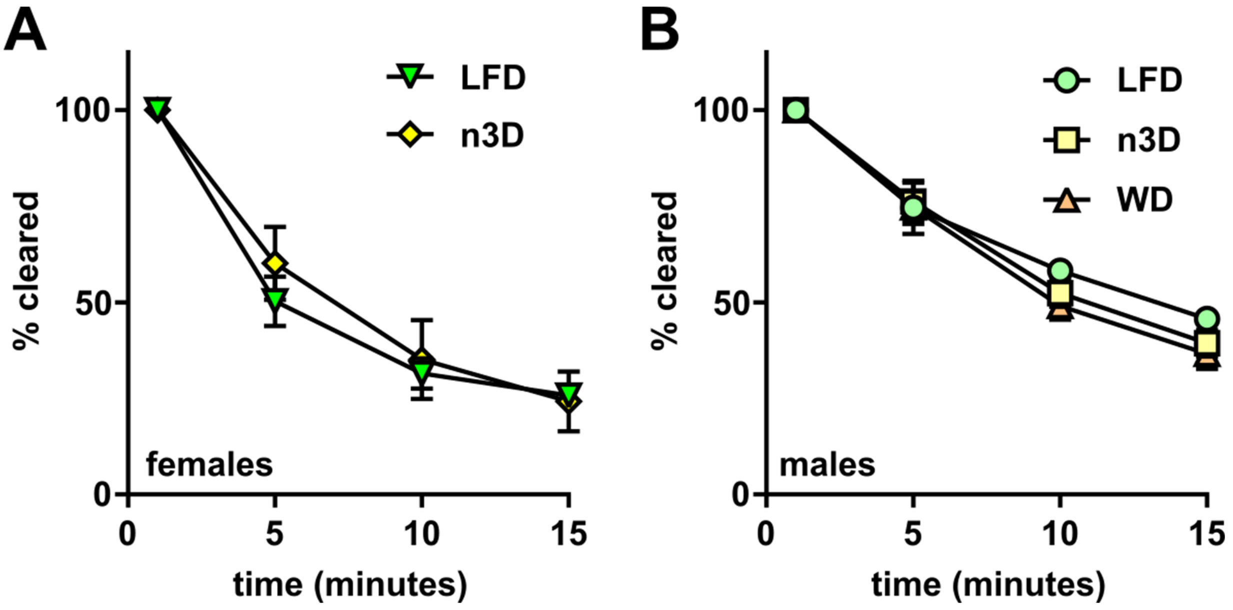


**Supplementary Figure 4: Plasma triglyceride clearance in 2-year old mice.** Mice were fasted (6 h) and injected intravenously with $^3$H-triglyceride containing chylomicrons. Blood was collected after 1, 5, 10, and 15 min after injection to measure clearance of radiolabel from the plasma. Points represent radiolabel remaining in the plasma as a percentage of the 1-min time point (means ± SEM) in 2-year-old female **(A)** and male **(B)** mice.

**Supplementary Figure 5**

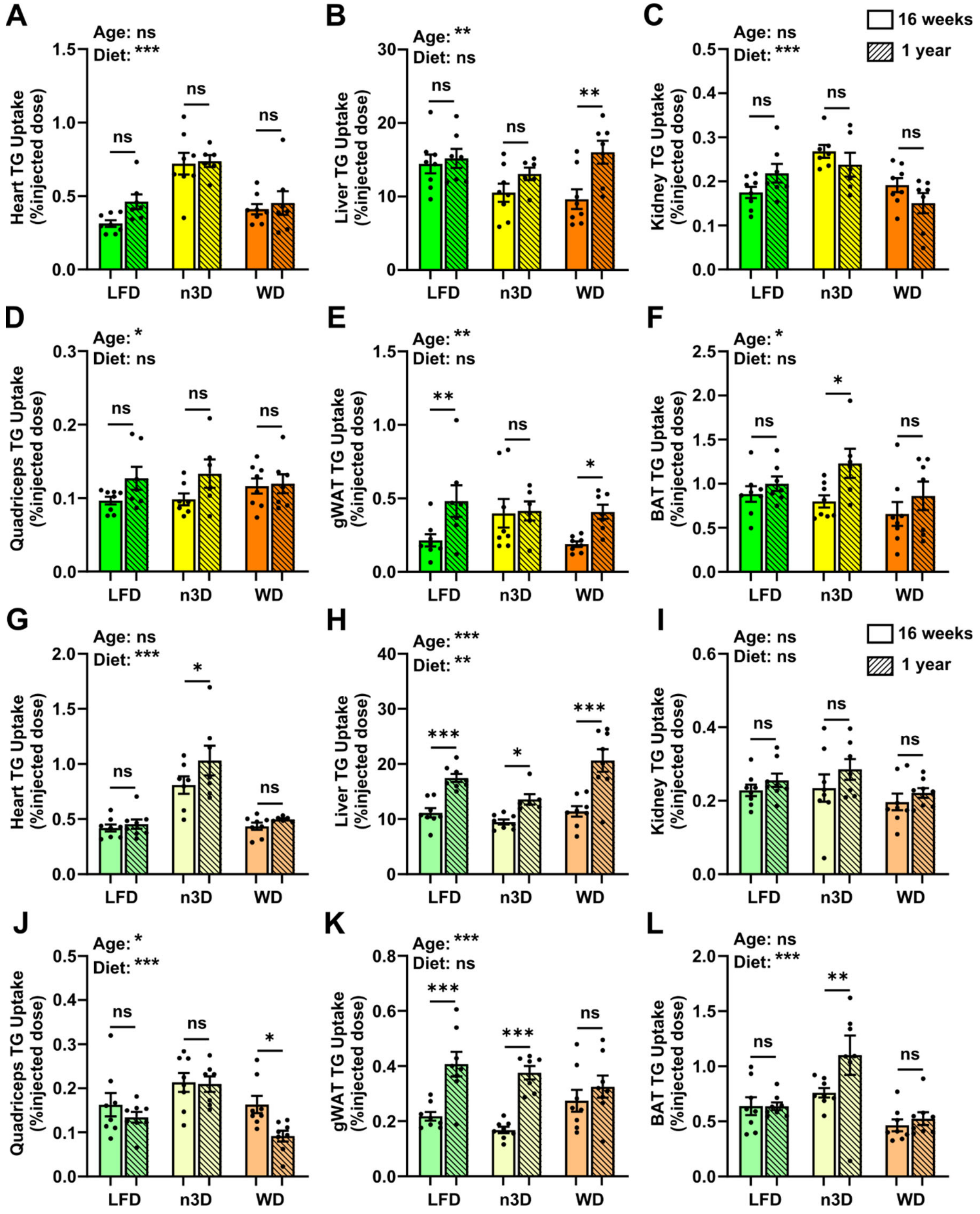


**Supplementary Figure 5: Total tissue triglyceride uptake.** At 16 weeks or 1 year of age female **(A-F)** and male **(G-L)** mice were fasted (6 h) and injected intravenously with $^3$H-triglyceride containing chylomicrons. After 15 minutes, tissues were harvested and uptake of radiolabel (% injected dose for total tissue) was measured in heart **(A,G)**, liver **(B,H)**, kidney **(C,I)**, quadriceps muscle **(D,J)**, gonadal white adipose tissue (gWAT)**(E,K)**, and brown adipose tissue (BAT)**(F,L)**. Bars represent mean ± SEM. Ns=not significant, *p<0.05, **p<0.01, ***p<0.001 by 2-way ANOVA and subsequent Tukey's multiple comparison test. Complete lists of p values are listed in Supplementary Table 1.

**Supplementary Figure 6**

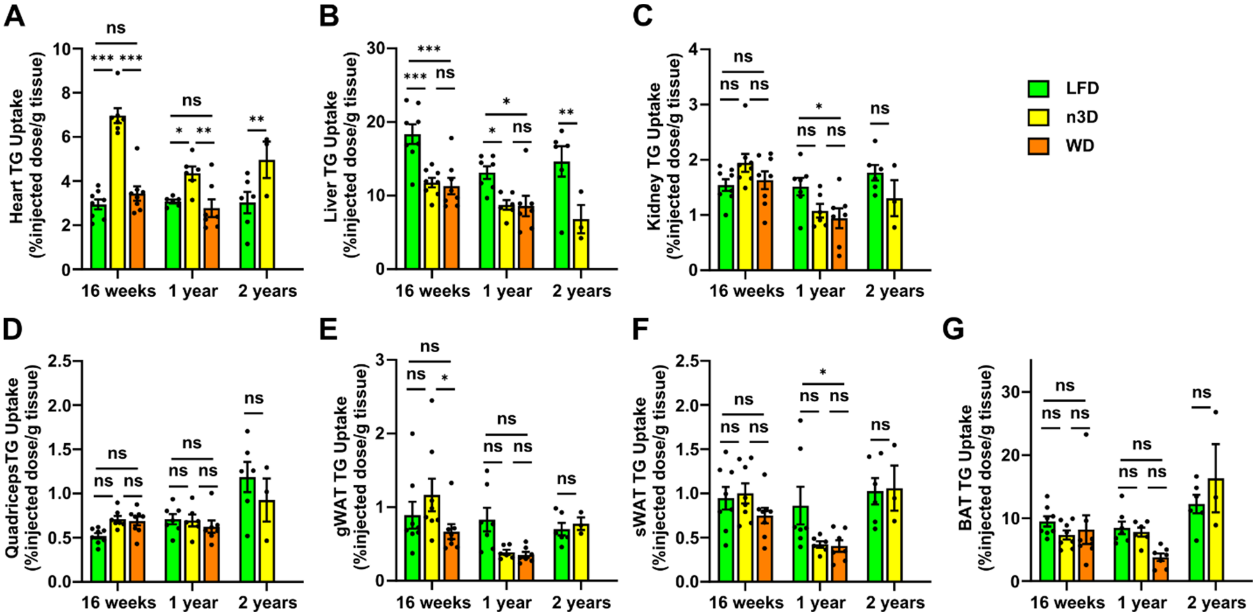


**Supplementary Figure 6: Tissue triglyceride uptake in female mice.** At 16 weeks, 1 year, or 2 years of age female mice were fasted (6 h) and injected intravenously with $^{3}$H-triglyceride containing chylomicrons. After 15 minutes, tissues were harvested and uptake of radiolabel (% injected dose/g tissue) was measured in heart **(A)**, liver **(B)**, kidney **(C)**, quadriceps muscle **(D)**, gonadal white adipose tissue (gWAT)**(E)**, subcutaneous adipose tissue (sWAT)**(F)**, and brown adipose tissue (BAT)**(G)**. Bars represent mean ± SEM. ns=not significant, *p<0.05, **p<0.01, ***p<0.001 by Tukey's multiple comparison test after 2-way ANOVA. Complete lists of p values are listed in Supplementary Table 1.

**Supplementary Figure 7**

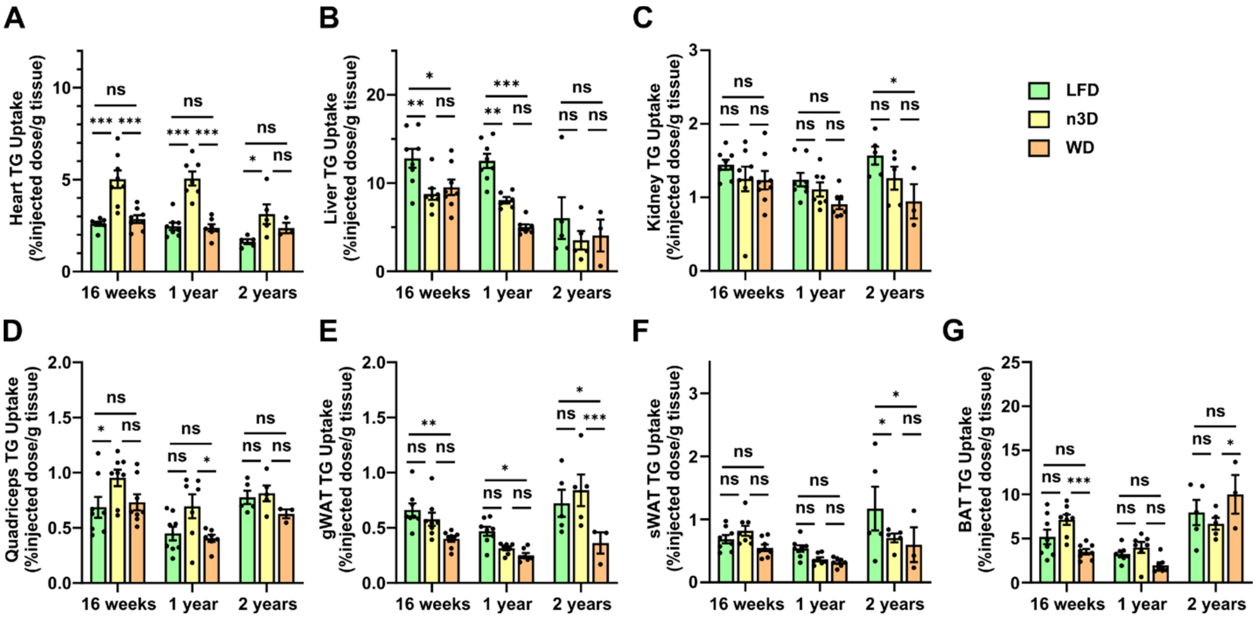


**Supplementary Figure 7: Tissue triglyceride uptake in male mice.** At 16 weeks, 1 year, or 2 years of age male mice were fasted (6 h) and injected intravenously with $^{3}$H-triglyceride containing chylomicrons. After 15 minutes, tissues were harvested and uptake of radiolabel (% injected dose/g tissue) was measured in heart **(A)**, liver **(B)**, kidney **(C)**, quadriceps muscle **(D)**, gonadal white adipose tissue (gWAT)**(E)**, subcutaneous adipose tissue (sWAT)**(F)**, and brown adipose tissue (BAT)**(G)**. Bars represent mean ± SEM. ns=not significant, *p<0.05, **p<0.01, ***p<0.001 by Tukey's multiple comparison test after 2-way ANOVA. Complete lists of p values are listed in Supplementary Table 1.

**Supplementary Figure 8**

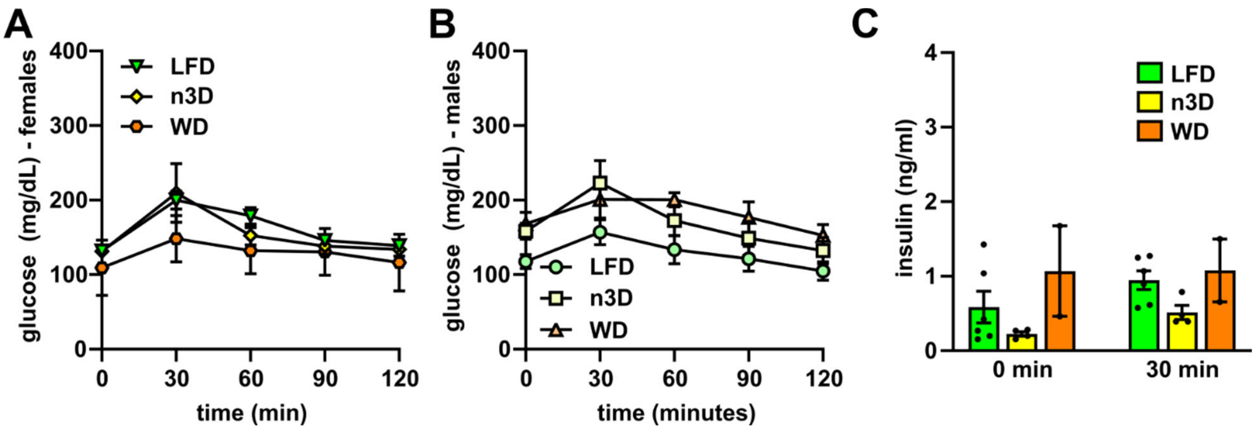


**Supplementary Figure 8: Glucose tolerance tests and plasma insulin levels in old mice.** **A-B)** Glucose tolerance tests were performed on fasted (6 h) female (A) and male (B) mice at 2 years of age. Mice were injected with glucose (1 g/kg) and blood glucose concentrations were measured over 2 h. **C)** Plasma insulin levels before and 30 minutes after glucose injection in 102-week-old female mice.

**Supplementary Figure 9**

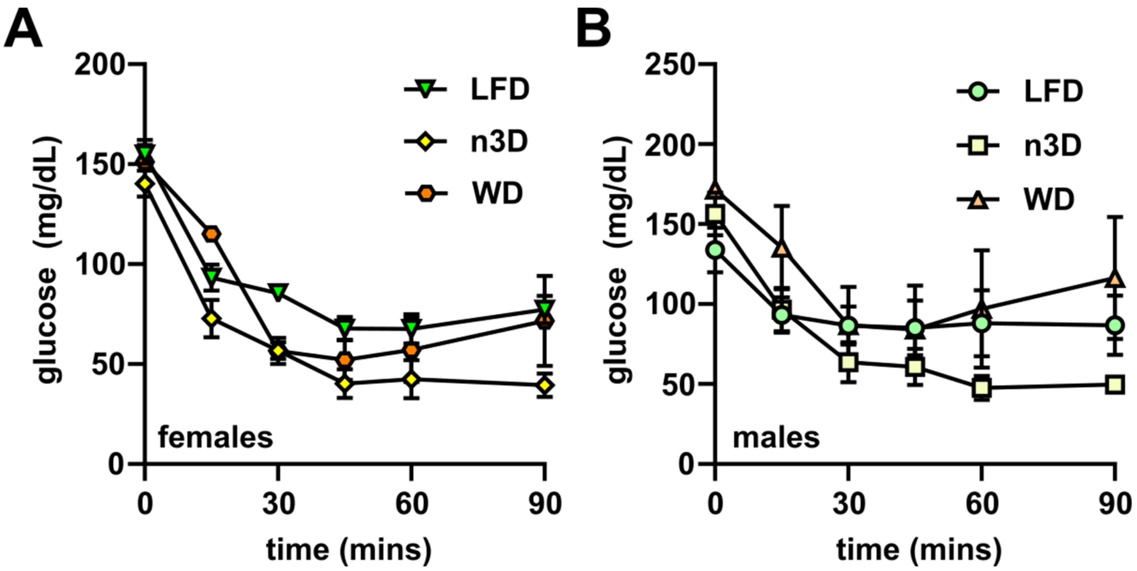


**Supplementary Figure 9: Insulin tolerance tests.** Insulin tolerance tests were performed in fasted (4 h) female **(A)** and male **(B)** mice at 2 years of age. Mice were injected with 0.5 U/mL of human insulin (Humalin-R) and blood glucose concentrations were measured over 90 min. Points represent glucose levels (means ± SEM) at each respective time point.

**Supplementary Table 1**

**Figure 1 statistics**

Panel B (All Survival)

| | # of animals |
|---|---|
| LFD | 18 |
| n3D | 18 |
| WD | 18 |
| Survival Log-Rank (Mantel-Cox) test | |
| | P value |
| LFD vs n3D | 0.409 |
| LFD vs WD | 0.003** |
| n3D vs WD | 0.054 |

Panel C (Female Survival)

| | # of animals |
|---|---|
| LFD | 10 |
| n3D | 10 |
| WD | 10 |
| Survival Log-Rank (Mantel-Cox) test | |
| | P value |
| LFD vs n3D | 0.400 |
| LFD vs WD | 0.013* |
| n3D vs WD | 0.116 |

Panel D (Male Survival)

| | # of animals |
|---|---|
| LFD | 8 |
| n3D | 8 |
| WD | 8 |
| Survival Log-Rank (Mantel-Cox) test | |
| | P value |
| LFD vs n3D | 0.861 |
| LFD vs WD | 0.149 |
| n3D vs WD | 0.236 |

**Figure 2 statistics**

Panel A (Female body weights)

| | # of animals |
|---|---|
| LFD | 10 at start |
| n3D | 10 at start |
| WD | 10 at start |

Panel B (Male body weights)

| | # of animals |
|---|---|
| LFD | 8 at start |
| n3D | 8 at start |
| WD | 8 at start |

Panel C (Female food consumption)

| | # of cages (animals) |
|---|---|
| LFD | 5 (17) |
| n3D | 5 (15) |
| WD | 7 (16) |
| | P value |
| 1-way ANOVA | 0.001** |
| Tukey's multiple comparison | |
| LFD vs n3D | 0.262 |
| LFD vs WD | 0.001** |
| n3D vs WD | 0.030* |

Panel D (Male food consumption)

| | # of cages (animals) |
|---|---|
| LFD | 4 (16) |
| n3D | 4 (15) |
| WD | 4 (16) |
| | P value |
| 1-way ANOVA | 0.003** |
| Tukey's multiple comparison | |
| LFD vs n3D | 0.915 |
| LFD vs WD | 0.009** |
| n3D vs WD | 0.003** |

**Figure 3 statistics**

Panel A (Female Lean Mass)

| # of animals | # of animals |
|---|---|
| LFD | |
| 8 weeks (pre-diet) | 26 |
| 16 weeks | 26 |
| 8 months | 16 (18-2 mice died=16) |
| 1 year | 15 (18-3 mice died=15) |
| 1.5 year | 8 (10-2 mice died=8) |
| n3D | |
| 8 weeks (pre-diet) | 25 (26-1 outlier value) |
| 16 weeks | 26 |
| 8 months | 15 (18-3 mice died=15) |
| 1 year | 14 (18-4 mice died=14) |
| 1.5 year | 7 (10-3 mice died=7) |
| WD | |
| 8 weeks (pre-diet) | 26 |
| 16 weeks | 26 |
| 8 months | 17 (18-1 mouse died=17 |
| 1 year | 15 (18-3 mice died=15) |
| 1.5 year | 5 (10-5 mice died=5) |
| | P value |
| 2-way ANOVA | |
| Age | <0.001*** |
| Diet | <0.001*** |
| Tukey's multiple comparison | |
| 8 weeks (pre-diet) | |

| | |
|---|---|
| LFD vs n3D | 0.687 |
| LFD vs WD | 0.998 |
| n3D vs WD | 0.721 |
| 16 weeks | |
| LFD vs n3D | <0.001*** |
| LFD vs WD | 0.003** |
| n3D vs WD | 0.774 |
| 8 months | |
| LFD vs n3D | <0.001*** |
| LFD vs WD | 0.004** |
| n3D vs WD | 0.002** |
| 1 year | |
| LFD vs n3D | <0.001*** |
| LFD vs WD | 0.002** |
| n3D vs WD | 0.018* |
| 1.5 years | |
| LFD vs n3D | <0.001*** |
| LFD vs WD | <0.001*** |
| n3D vs WD | 0.469 |
| LFD | |
| 8 weeks (pre-diet) vs. 16 weeks | 0.007** |
| 8 weeks (pre-diet) vs. 8 months | <0.001*** |
| 8 weeks (pre-diet) vs. 1 year | <0.001*** |
| 8 weeks (pre-diet) vs. 1.5 years | <0.001*** |
| 16 weeks vs. 8 months | <0.001*** |
| 16 weeks vs. 1 year | <0.001*** |
| 16 weeks vs. 1.5 years | <0.001*** |
| 8 months vs. 1 year | 0.253 |
| 8 months vs. 1.5 years | 0.521 |
| 1 year vs. 1.5 years | >0.999 |
| n3D | |
| 8 weeks (pre-diet) vs. 16 weeks | <0.001*** |
| 8 weeks (pre-diet) vs. 8 months | <0.001*** |
| 8 weeks (pre-diet) vs. 1 year | <0.001*** |
| 8 weeks (pre-diet) vs. 1.5 years | <0.001*** |
| 16 weeks vs. 8 months | <0.001*** |
| 16 weeks vs. 1 year | <0.001*** |
| 16 weeks vs. 1.5 years | <0.001*** |
| 8 months vs. 1 year | 0.372 |
| 8 months vs. 1.5 years | 0.061 |
| 1 year vs. 1.5 years | 0.756 |
| WD | |
| 8 weeks (pre-diet) vs. 16 weeks | <0.001*** |
| 8 weeks (pre-diet) vs. 8 months | <0.001*** |
| 8 weeks (pre-diet) vs. 1 year | <0.001*** |
| 8 weeks (pre-diet) vs. 1.5 years | <0.001*** |
| 16 weeks vs. 8 months | <0.001*** |
| 16 weeks vs. 1 year | <0.001*** |
| 16 weeks vs. 1.5 years | <0.001*** |
| 8 months vs. 1 year | 0.096 |
| 8 months vs. 1.5 years | 0.006** |
| 1 year vs. 1.5 years | 0.430 |

Panel B (Male Lean Mass)

| | # of animals |
|---|---|
| LFD | |
| 8 weeks (pre-diet) | 24 |
| 16 weeks | 24 |
| 8 months | 16 |
| 1 year | 16 |
| 1.5 year | 8 |
| n3D | |
| 8 weeks (pre-diet) | 24 |
| 16 weeks | 24 |
| 8 months | 14 (16-1 mouse died-1 mouse not measured=14) |
| 1 year | 15 (16-1 mouse died) |
| 1.5 year | 7 (8-1 mouse died=7) |
| WD | |
| 8 weeks (pre-diet) | 24 |
| 16 weeks | 24 |
| 8 months | 16 |
| 1 year | 14 (16-2 mice not measured=14) |
| 1.5 year | 5 (8-3 mice died=5) |
| | P value |
| 2-way ANOVA | |
| Age | <0.001*** |
| Diet | <0.001*** |
| Tukey's multiple comparison | |
| 8 weeks (pre-diet) | |
| LFD vs n3D | 0.648 |
| LFD vs WD | 0.946 |
| n3D vs WD | 0.451 |
| 16 weeks | |
| LFD vs n3D | 0.464 |
| LFD vs WD | 0.044* |
| n3D vs WD | 0.439 |
| 8 months | |
| LFD vs n3D | 0.182 |
| LFD vs WD | <0.001*** |
| n3D vs WD | 0.017* |
| 1 year | |
| LFD vs n3D | 0.006** |
| LFD vs WD | <0.001*** |
| n3D vs WD | 0.008** |
| 1.5 years | |
| LFD vs n3D | <0.001*** |
| LFD vs WD | 0.003** |
| n3D vs WD | 0.917 |
| LFD | |
| 8 weeks (pre-diet) vs. 16 weeks | <0.001*** |
| 8 weeks (pre-diet) vs. 8 months | <0.001*** |
| 8 weeks (pre-diet) vs. 1 year | <0.001*** |
| 8 weeks (pre-diet) vs. 1.5 years | <0.001*** |
| 16 weeks vs. 8 months | 0.007** |
| 16 weeks vs. 1 year | <0.001*** |

| | |
|---|---|
| 16 weeks vs. 1.5 years | 0.047* |
| 8 months vs. 1 year | 0.974 |
| 8 months vs. 1.5 years | >0.999 |
| 1 year vs. 1.5 years | 0.993 |
| n3D | |
| 8 weeks (pre-diet) vs. 16 weeks | <0.001*** |
| 8 weeks (pre-diet) vs. 8 months | <0.001*** |
| 8 weeks (pre-diet) vs. 1 year | <0.001*** |
| 8 weeks (pre-diet) vs. 1.5 years | <0.001*** |
| 16 weeks vs. 8 months | <0.001*** |
| 16 weeks vs. 1 year | <0.001*** |
| 16 weeks vs. 1.5 years | <0.001*** |
| 8 months vs. 1 year | 0.363 |
| 8 months vs. 1.5 years | 0.013* |
| 1 year vs. 1.5 years | 0.396 |
| WD | |
| 8 weeks (pre-diet) vs. 16 weeks | <0.001*** |
| 8 weeks (pre-diet) vs. 8 months | <0.001*** |
| 8 weeks (pre-diet) vs. 1 year | <0.001*** |
| 8 weeks (pre-diet) vs. 1.5 years | <0.001*** |
| 16 weeks vs. 8 months | <0.001*** |
| 16 weeks vs. 1 year | <0.001*** |
| 16 weeks vs. 1.5 years | <0.001*** |
| 8 months vs. 1 year | 0.203 |
| 8 months vs. 1.5 years | 0.990 |
| 1 year vs. 1.5 years | 0.832 |

Panel C (Female Fat Mass)

| | # of animals |
|---|---|
| LFD | |
| 8 weeks (pre-diet) | 26 |
| 16 weeks | 26 |
| 8 months | 16 (18-2 mice died=16) |
| 1 year | 15 (18-3 mice died=15) |
| 1.5 year | 8 (10-2 mice died=8) |
| n3D | |
| 8 weeks (pre-diet) | 26 |
| 16 weeks | 26 |
| 8 months | 15 (18-3 mice died=15) |
| 1 year | 14 (18-4 mice died=14) |
| 1.5 year | 7 (10-3 mice died=7) |
| WD | |
| 8 weeks (pre-diet) | 26 |
| 16 weeks | 24 (26-2 outlier values) |
| 8 months | 17 (18-1 mouse died=17) |
| 1 year | 15 (18-3 mice died=15) |
| 1.5 year | 5 (10-5 mice died=5) |
| | P value |
| 2-way ANOVA | |
| Age | <0.001*** |
| Diet | <0.001*** |
| Tukey’s multiple comparison | |
| 8 weeks (pre-diet) | |

| | |
|---|---|
| LFD vs n3D | 0.972 |
| LFD vs WD | 0.992 |
| n3D vs WD | 0.994 |
| 16 weeks | |
| LFD vs n3D | 0.208 |
| LFD vs WD | 0.934 |
| n3D vs WD | 0.390 |
| 8 months | |
| LFD vs n3D | 0.472 |
| LFD vs WD | <0.001*** |
| n3D vs WD | <0.001*** |
| 1 year | |
| LFD vs n3D | 0.377 |
| LFD vs WD | <0.001*** |
| n3D vs WD | <0.001*** |
| 1.5 years | |
| LFD vs n3D | 0.997 |
| LFD vs WD | <0.001*** |
| n3D vs WD | <0.001*** |
| LFD | |
| 8 weeks (pre-diet) vs. 16 weeks | <0.001*** |
| 8 weeks (pre-diet) vs. 8 months | <0.001*** |
| 8 weeks (pre-diet) vs. 1 year | <0.001*** |
| 8 weeks (pre-diet) vs. 1.5 years | <0.001*** |
| 16 weeks vs. 8 months | 0.930 |
| 16 weeks vs. 1 year | <0.001*** |
| 16 weeks vs. 1.5 years | <0.001*** |
| 8 months vs. 1 year | 0.002** |
| 8 months vs. 1.5 years | <0.001*** |
| 1 year vs. 1.5 years | 0.891 |
| n3D | |
| 8 weeks (pre-diet) vs. 16 weeks | 0.015* |
| 8 weeks (pre-diet) vs. 8 months | <0.001*** |
| 8 weeks (pre-diet) vs. 1 year | <0.001*** |
| 8 weeks (pre-diet) vs. 1.5 years | <0.001*** |
| 16 weeks vs. 8 months | 0.004** |
| 16 weeks vs. 1 year | <0.001*** |
| 16 weeks vs. 1.5 years | <0.001*** |
| 8 months vs. 1 year | 0.001** |
| 8 months vs. 1.5 years | 0.040* |
| 1 year vs. 1.5 years | 0.999 |
| WD | |
| 8 weeks (pre-diet) vs. 16 weeks | <0.001*** |
| 8 weeks (pre-diet) vs. 8 months | <0.001*** |
| 8 weeks (pre-diet) vs. 1 year | <0.001*** |
| 8 weeks (pre-diet) vs. 1.5 years | <0.001*** |
| 16 weeks vs. 8 months | <0.001*** |
| 16 weeks vs. 1 year | <0.001*** |
| 16 weeks vs. 1.5 years | <0.001*** |
| 8 months vs. 1 year | <0.001*** |
| 8 months vs. 1.5 years | <0.001*** |
| 1 year vs. 1.5 years | <0.001*** |

Panel D (Male Fat Mass)

| | # of animals |
|---|---|
| LFD | |
| 8 weeks (pre-diet) | 24 |
| 16 weeks | 23 (24-1 outlier value=23) |
| 8 months | 16 |
| 1 year | 16 |
| 1.5 year | 8 |
| n3D | |
| 8 weeks (pre-diet) | 24 |
| 16 weeks | 24 |
| 8 months | 14 (16-1 mouse died-1 mouse not measured=14) |
| 1 year | 15 (16-1 mouse died) |
| 1.5 year | 7 (8-1 mouse died=7) |
| WD | |
| 8 weeks (pre-diet) | 24 |
| 16 weeks | 24 |
| 8 months | 16 |
| 1 year | 14 (16-2 mice not measured) |
| 1.5 year | 5 (8-3 mice died=5) |
| | P value |
| 2-way ANOVA | |
| Age | <0.001*** |
| Diet | <0.001*** |
| Tukey's multiple comparison | |
| 8 weeks (pre-diet) | |
| LFD vs n3D | 0.995 |
| LFD vs WD | 0.996 |
| n3D vs WD | 0.999 |
| 16 weeks | |
| LFD vs n3D | 0.345 |
| LFD vs WD | <0.001*** |
| n3D vs WD | <0.001*** |
| 8 months | |
| LFD vs n3D | 0.013* |
| LFD vs WD | <0.001*** |
| n3D vs WD | <0.001*** |
| 1 year | |
| LFD vs n3D | 0.539 |
| LFD vs WD | <0.001*** |
| n3D vs WD | <0.001*** |
| 1.5 years | |
| LFD vs n3D | 0.008** |
| LFD vs WD | <0.001*** |
| n3D vs WD | 0.115 |
| LFD | |
| 8 weeks (pre-diet) vs. 16 weeks | 0.022* |
| 8 weeks (pre-diet) vs. 8 months | <0.001*** |
| 8 weeks (pre-diet) vs. 1 year | <0.001*** |
| 8 weeks (pre-diet) vs. 1.5 years | <0.001*** |
| 16 weeks vs. 8 months | <0.001*** |
| 16 weeks vs. 1 year | <0.001*** |

| | |
|---|---|
| 16 weeks vs. 1.5 years | 0.035* |
| 8 months vs. 1 year | 0.099 |
| 8 months vs. 1.5 years | 0.947 |
| 1 year vs. 1.5 years | 0.048* |
| n3D | |
| 8 weeks (pre-diet) vs. 16 weeks | 0.401 |
| 8 weeks (pre-diet) vs. 8 months | <0.001*** |
| 8 weeks (pre-diet) vs. 1 year | <0.001*** |
| 8 weeks (pre-diet) vs. 1.5 years | <0.001*** |
| 16 weeks vs. 8 months | <0.001*** |
| 16 weeks vs. 1 year | <0.001*** |
| 16 weeks vs. 1.5 years | <0.001*** |
| 8 months vs. 1 year | 0.980 |
| 8 months vs. 1.5 years | 0.994 |
| 1 year vs. 1.5 years | >0.999 |
| WD | |
| 8 weeks (pre-diet) vs. 16 weeks | <0.001*** |
| 8 weeks (pre-diet) vs. 8 months | <0.001*** |
| 8 weeks (pre-diet) vs. 1 year | <0.001*** |
| 8 weeks (pre-diet) vs. 1.5 years | <0.001*** |
| 16 weeks vs. 8 months | <0.001*** |
| 16 weeks vs. 1 year | <0.001*** |
| 16 weeks vs. 1.5 years | <0.001*** |
| 8 months vs. 1 year | <0.001*** |
| 8 months vs. 1.5 years | 0.670 |
| 1 year vs. 1.5 years | <0.001*** |

**Figure 4 statistics**

Panel A (Plasma TG females)

| | # of animals |
|---|---|
| LFD | |
| 8 weeks (pre-diet) | 10 |
| 16 weeks | 10 |
| 8 months | 9 (10-1 mouse died=9) |
| 1 year | 8 (10-2 mice died=8) |
| 1.5 year | 8 (10-2 mice died=8) |
| n3D | |
| 8 weeks (pre-diet) | 10 |
| 16 weeks | 10 |
| 8 months | 9 (10-1 mouse died=9) |
| 1 year | 8 (10-2 mice died=8) |
| 1.5 year | 8 (10-2 mice died=8) |
| WD | |
| 8 weeks (pre-diet) | 10 |
| 16 weeks | 10 |
| 8 months | 10 |
| 1 year | 7 (10-2 mice died-1 outlier value=7) |
| 1.5 year | 5 (10-5 mice died=5) |
| | P value |
| Mixed effects analysis | |
| Age | <0.001*** |

| | |
|---|---|
| Diet | 0.001** |
| Tukey's multiple comparison | |
| 8 weeks (pre-diet) | |
| LFD vs n3D | 0.340 |
| LFD vs WD | 0.370 |
| n3D vs WD | 0.941 |
| 16 weeks | |
| LFD vs n3D | 0.055 |
| LFD vs WD | 0.977 |
| n3D vs WD | 0.106 |
| 8 months | |
| LFD vs n3D | 0.449 |
| LFD vs WD | 0.940 |
| n3D vs WD | 0.358 |
| 1 year | |
| LFD vs n3D | 0.194 |
| LFD vs WD | 0.324 |
| n3D vs WD | 0.683 |
| 1.5 years | |
| LFD vs n3D | 0.004** |
| LFD vs WD | 0.953 |
| n3D vs WD | 0.126 |

Panel B (Plasma TG males)

| | # of animals |
|---|---|
| LFD | |
| 8 weeks (pre-diet) | 8 |
| 16 weeks | 8 |
| 8 months | 8 |
| 1 year | 8 |
| 1.5 year | 8 |
| n3D | |
| 8 weeks (pre-diet) | 8 |
| 16 weeks | 8 |
| 8 months | 7 (8-1 sample lost=7) |
| 1 year | 8 |
| 1.5 year | 7 (8-1 mouse died=7) |
| WD | |
| 8 weeks (pre-diet) | 8 |
| 16 weeks | 8 |
| 8 months | 8 |
| 1 year | 7 (8-1 sample lost=7) |
| 1.5 year | 5 (8-3 mice died=5) |
| | P value |
| Mixed effects analysis | |
| Age | <0.001*** |
| Diet | 0.088 |
| Tukey's multiple comparison | |
| 8 weeks (pre-diet) | |
| LFD vs n3D | 0.804 |
| LFD vs WD | 0.604 |
| n3D vs WD | 0.832 |
| 16 weeks | |

| | |
|---|---|
| LFD vs n3D | 0.086 |
| LFD vs WD | 0.999 |
| n3D vs WD | 0.124 |
| 8 months | |
| LFD vs n3D | 0.326 |
| LFD vs WD | 0.936 |
| n3D vs WD | 0.254 |
| 1 year | |
| LFD vs n3D | 0.961 |
| LFD vs WD | 0.137 |
| n3D vs WD | 0.397 |
| 1.5 years | |
| LFD vs n3D | 0.809 |
| LFD vs WD | 0.441 |
| n3D vs WD | 0.572 |

**Figure 5 statistics**

Panel A (Female FTT 13 weeks)

| | # of animals |
|---|---|
| LFD | 8 |
| n3D | 8 |
| WD | 8 |
| | P-Value |
| 1-Way ANOVA of AUC | 0.047* |
| Tukey's multiple comparison | |
| LFD vs n3D | 0.039* |
| LFD vs WD | 0.584 |
| n3D vs WD | 0.252 |

Panel B (Male FTT 13 weeks)

| | # of animals |
|---|---|
| LFD | 8 |
| n3D | 8 |
| WD | 8 |
| | P-Value |
| 1-Way ANOVA of AUC | 0.778 |
| Tukey's multiple comparison | |
| LFD vs n3D | 0.900 |
| LFD vs WD | 0.763 |
| n3D vs WD | 0.961 |

Panel C (Female FTT 1 year)

| # of animals | # of animals |
|---|---|
| LFD | 7 (8-1 mouse died=7) |
| n3D | 6 (8-2 mice died=6) |
| WD | 7 (8-1 mouse died=7) |
| | P-Value |
| 1-Way ANOVA of AUC | <0.001*** |
| Tukey's multiple comparison | |
| LFD vs n3D | <0.001*** |
| LFD vs WD | 0.961 |
| n3D vs WD | <0.001*** |

Panel D (Male FTT 1 year)

| | # of animals |
|---|---|
| LFD | 7 (8-1 outlier value=7) |
| n3D | 7 (8-1 mouse died=7) |
| WD | 8 |
| | P-Value |
| 1-Way ANOVA of AUC | 0.013* |
| Tukey's multiple comparison | |
| LFD vs n3D | 0.020* |
| LFD vs WD | 0.954 |
| n3D vs WD | 0.030* |

Panel E (Female FTT 1.5 year)

| | # of animals |
|---|---|
| LFD | 8 (10-2 mice died=8) |
| n3D | 8 (10-2 mice died=8) |
| WD | 4 (10-5 mice died-1 outlier value=4) |
| | P-Value |
| 1-Way ANOVA of AUC | 0.001** |
| Tukey's multiple comparison | |
| LFD vs n3D | 0.010** |
| LFD vs WD | 0.384 |
| n3D vs WD | 0.002** |

Panel F (Male FTT 1.5 year)

| | # of animals |
|---|---|
| LFD | 8 |
| n3D | 7 (8-1 mouse died=7) |
| WD | 5 (8-3 mice died=5) |
| | P-Value |
| 1-Way ANOVA of AUC | 0.054 |
| Tukey's multiple comparison | |
| LFD vs n3D | 0.107 |
| LFD vs WD | 0.891 |
| n3D vs WD | 0.074 |

**Figure 6 statistics**

Panel A (Female 16-week chylomicron clearance)

| | # of animals |
|---|---|
| LFD | 8 |
| n3D | 8 |
| WD | 8 |
| | P value |
| 2-way repeated measures ANOVA | |
| Diet | 0.1332 |
| Time | <0.001*** |
| Tukey's multiple comparison | |
| 5 minutes | |
| LFD vs n3D | 0.007** |
| LFD vs WD | 0.571 |
| n3D vs WD | 0.310 |

| | |
|---|---|
| 10 minutes | |
| LFD vs n3D | 0.169 |
| LFD vs WD | 0.866 |
| n3D vs WD | 0.625 |
| 15 minutes | |
| LFD vs n3D | 0.995 |
| LFD vs WD | 0.665 |
| n3D vs WD | 0.832 |

Panel B (Male 16-week chylomicron clearance)

| | # of animals |
|---|---|
| LFD | 8 |
| n3D | 8 |
| WD | 8 |
| | P value |
| 2-way repeated measures ANOVA | |
| Diet | 0.418 |
| Time | <0.001*** |
| Tukey's multiple comparison | |
| 5 minutes | |
| LFD vs n3D | 0.648 |
| LFD vs WD | 0.927 |
| n3D vs WD | 0.523 |
| 10 minutes | |
| LFD vs n3D | 0.522 |
| LFD vs WD | 0.778 |
| n3D vs WD | 0.341 |
| 15 minutes | |
| LFD vs n3D | 0.884 |
| LFD vs WD | 0.250 |
| n3D vs WD | 0.670 |

Panel C (Female 1-year chylomicron clearance)

| | # of animals |
|---|---|
| LFD | 7 (8-1 mouse died=7) |
| n3D | 6 (8-2 mice died=6) |
| WD | 7 (8-1 mouse died=7) |
| | P value |
| 2-way repeated measures ANOVA | |
| Diet | 0.444 |
| Time | <0.001** |
| Tukey's multiple comparison | |
| 5 minutes | |
| LFD vs n3D | 0.946 |
| LFD vs WD | 0.799 |
| n3D vs WD | 0.519 |
| 10 minutes | |
| LFD vs n3D | 0.521 |
| LFD vs WD | 0.428 |
| n3D vs WD | 0.921 |
| 15 minutes | |
| LFD vs n3D | 0.076 |
| LFD vs WD | 0.140 |
| n3D vs WD | 0.983 |

Panel D (Male 1-year chylomicron clearance)

| | # of animals |
|---|---|
| LFD | 8 |
| n3D | 7 (8-1 mouse died=7) |
| WD | 7 (8-1 mouse died=7) |
| | P value |
| 2-way repeated measures ANOVA | |
| Diet | <0.001** |
| Time | <0.001** |
| Tukey's multiple comparison | |
| 5 minutes | |
| LFD vs n3D | 0.361 |
| LFD vs WD | 0.062 |
| n3D vs WD | 0.014* |
| 10 minutes | |
| LFD vs n3D | 0.269 |
| LFD vs WD | 0.035* |
| n3D vs WD | 0.005** |
| 15 minutes | |
| LFD vs n3D | 0.576 |
| LFD vs WD | <0.001** |
| n3D vs WD | 0.003** |

**<u>Figure 7 statistics</u>**

Panel A (Female Heart TG uptake)

| | # of animals |
|---|---|
| LFD | |
| 16 weeks | 8 |
| 1 year | 6 (8-1 mouse died-1 outlier value=6) |
| n3D | |
| 16 weeks | 7 (8-1 outlier value=7) |
| 1 year | 6 (8-2 mice died=6) |
| WD | |
| 16 weeks | 8 |
| 1 year | 7 (8-1 mouse died=7) |
| | P value |
| 2-way ANOVA | |
| Age | <0.001*** |
| Diet | <0.001*** |
| Tukey's multiple comparison | |
| 16 weeks | |
| LFD vs n3D | <0.001*** |
| LFD vs WD | 0.445 |
| n3D vs WD | <0.001*** |
| 1 year | |
| LFD vs n3D | 0.024* |
| LFD vs WD | 0.786 |
| n3D vs WD | 0.003** |
| LFD | |
| 16 weeks vs. 1 year | 0.764 |
| n3D | |

| | |
|---|---|
| 16 weeks vs. 1 year | <0.001*** |
| WD | |
| 16 weeks vs. 1 year | 0.122 |

Panel B (Female Liver TG uptake)

| | # of animals |
|---|---|
| LFD | |
| 16 weeks | 8 |
| 1 year | 7 (8-1 mouse died=7) |
| n3D | |
| 16 weeks | 8 |
| 1 year | 6 (8-2 mice died=6) |
| WD | |
| 16 weeks | 8 |
| 1 year | 7 (8-1 mouse died=7) |
| | P value |
| 2-way ANOVA | |
| Age | <0.001*** |
| Diet | <0.001*** |
| Tukey's multiple comparison | |
| 16 weeks | |
| LFD vs n3D | <0.001*** |
| LFD vs WD | <0.001*** |
| n3D vs WD | 0.942 |
| 1 year | |
| LFD vs n3D | 0.025* |
| LFD vs WD | 0.015* |
| n3D vs WD | 0.995 |
| LFD | |
| 16 weeks vs. 1 year | 0.001** |
| n3D | |
| 16 weeks vs. 1 year | 0.060 |
| WD | |
| 16 weeks vs. 1 year | 0.078 |

Panel C (Female Kidney TG uptake)

| | # of animals |
|---|---|
| LFD | |
| 16 weeks | 7 (8-1 mouse died=7) |
| 1 year | |
| n3D | 8 |
| 16 weeks | 6 (8-2 mice died=6) |
| 1 year | |
| WD | 8 |
| 16 weeks | 7 (8-1 mouse died=7) |
| 1 year | 7 (8-1 mouse died=7) |
| | P value |
| 2-way ANOVA | |
| Age | <0.001*** |
| Diet | 0.215 |
| Tukey's multiple comparison | |
| 16 weeks | |
| LFD vs n3D | 0.136 |
| LFD vs WD | 0.902 |

| | |
|---|---|
| n3D vs WD | 0.289 |
| 1 year | |
| LFD vs n3D | 0.145 |
| LFD vs WD | 0.033* |
| n3D vs WD | 0.827 |
| LFD | |
| 16 weeks vs. 1 year | 0.903 |
| n3D | |
| 16 weeks vs. 1 year | <0.001*** |
| WD | |
| 16 weeks vs. 1 year | 0.002** |

Panel D (Female Quadriceps TG uptake)

| | # of animals |
|---|---|
| LFD | |
| 16 weeks | 8 |
| 1 year | 7 (8-1 mouse died=7) |
| n3D | |
| 16 weeks | 7 (8-1 outlier value=7) |
| 1 year | 6 (8-2 mice died=6) |
| WD | |
| 16 weeks | 8 |
| 1 year | 7 (8-1 mouse died=7) |
| | P value |
| 2-way ANOVA | |
| Age | 0.395 |
| Diet | 0.273 |
| Tukey's multiple comparison | |
| 16 weeks | |
| LFD vs ω3FD | 0.036* |
| LFD vs WD | 0.055 |
| ω3FD vs WD | 0.960 |
| 1 year | |
| LFD vs ω3FD | 0.985 |
| LFD vs WD | 0.508 |
| ω3FD vs WD | 0.638 |
| LFD 16 weeks vs 1 year | 0.014* |
| ω3FD 16 weeks vs 1 year | 0.869 |
| WD 16 weeks vs 1 year | 0.383 |

Panel E (Female gWAT TG uptake)

| | # of animals |
|---|---|
| LFD | |
| 16 weeks | 8 |
| 1 year | 7 (8-1 mouse died=7) |
| n3D | |
| 16 weeks | 8 |
| 1 year | 6 (8-2 mice died=6) |
| WD | |
| 16 weeks | 8 |
| 1 year | 7 (8-1 mouse died=7) |
| | P value |
| 2-way ANOVA | |
| Age | 0.004** |

| | |
|---|---|
| Diet | 0.061 |
| Tukey's multiple comparison | |
| 16 weeks | |
| LFD vs ω3FD | 0.383 |
| LFD vs WD | 0.520 |
| ω3FD vs WD | 0.050* |
| 1 year | |
| LFD vs ω3FD | 0.137 |
| LFD vs WD | 0.084 |
| ω3FD vs WD | 0.987 |
| LFD 16 weeks vs 1 year | 0.774 |
| ω3FD 16 weeks vs 1 year | 0.001** |
| WD 16 weeks vs 1 year | 0.142 |

Panel F (Female sWAT TG uptake)

| | # of animals |
|---|---|
| LFD | |
| 16 weeks | 8 |
| 1 year | 7 (8-1 mouse died=7) |
| n3D | |
| 16 weeks | 8 |
| 1 year | 6 (8-2 mice died=6) |
| WD | |
| 16 weeks | 8 |
| 1 year | 7 (8-1 mouse died=7) |
| | P value |
| 2-way ANOVA | |
| Age | 0.002** |
| Diet | 0.035* |
| Tukey's multiple comparison | |
| 16 weeks | |
| LFD vs ω3FD | 0.946 |
| LFD vs WD | 0.467 |
| ω3FD vs WD | 0.296 |
| 1 year | |
| LFD vs ω3FD | 0.058 |
| LFD vs WD | 0.037* |
| ω3FD vs WD | 0.995 |
| LFD 16 weeks vs 1 year | 0.630 |
| ω3FD 16 weeks vs 1 year | 0.003** |
| WD 16 weeks vs 1 year | 0.053 |

Panel G (Female BAT TG uptake)

| | # of animals |
|---|---|
| LFD | |
| 16 weeks | 8 |
| 1 year | 7 (8-1 mouse died=7) |
| n3D | |
| 16 weeks | 8 |
| 1 year | 6 (8-2 mice died=6) |
| WD | |
| 16 weeks | 8 |
| 1 year | 7 (8-1 mouse died=7) |
| | P value |

| | |
|---|---|
| 2-way ANOVA | |
| Age | 0.112 |
| Diet | 0.064 |
| Tukey's multiple comparison | |
| 16 weeks | |
| LFD vs ω3FD | 0.415 |
| LFD vs WD | 0.722 |
| ω3FD vs WD | 0.868 |
| 1 year | |
| LFD vs ω3FD | 0.935 |
| LFD vs WD | 0.034* |
| ω3FD vs WD | 0.092 |
| LFD 16 weeks vs 1 year | 0.557 |
| ω3FD 16 weeks vs 1 year | 0.798 |
| WD 16 weeks vs 1 year | 0.015* |

**Figure 8 statistics**

Panel A (Male Heart TG uptake)

| | # of animals |
|---|---|
| LFD | |
| 16 weeks | 7 (8-outlier value=7) |
| 1 year | 8 |
| n3D | |
| 16 weeks | 8 |
| 1 year | 7 (8-1 mouse died=7) |
| WD | |
| 16 weeks | 8 |
| 1 year | 7 (8-1 mouse died=7) |
| | P value |
| 2-way ANOVA | |
| Age | 0.445 |
| Diet | <0.001*** |
| Tukey's multiple comparison | |
| 16 weeks | |
| LFD vs ω3FD | <0.001*** |
| LFD vs WD | 0.826 |
| ω3FD vs WD | <0.001*** |
| 1 year | |
| LFD vs ω3FD | <0.001*** |
| LFD vs WD | 0.979 |
| ω3FD vs WD | <0.001*** |
| LFD 16 weeks vs 1 year | 0.745 |
| ω3FD 16 weeks vs 1 year | 0.914 |
| WD 16 weeks vs 1 year | 0.271 |

Panel B (Male Liver TG uptake)

| | # of animals |
|---|---|
| LFD | |
| 16 weeks | 8 |
| 1 year | 8 |
| n3D | |
| 16 weeks | 8 |
| 1 year | 6 (8-1 mouse died-1 outlier value=6) |

| | |
|---|---|
| WD | |
| 16 weeks | 8 |
| 1 year | 7 (8-1 mouse died=7) |
| | P value |
| 2-way ANOVA | |
| Age | 0.007** |
| Diet | <0.001*** |
| Tukey's multiple comparison | |
| 16 weeks | |
| LFD vs ω3FD | 0.001** |
| LFD vs WD | 0.010* |
| ω3FD vs WD | 0.751 |
| 1 year | |
| LFD vs ω3FD | 0.001** |
| LFD vs WD | <0.001*** |
| ω3FD vs WD | 0.036* |
| LFD 16 weeks vs 1 year | 0.787 |
| ω3FD 16 weeks vs 1 year | 0.550 |
| WD 16 weeks vs 1 year | <0.001*** |

Panel C (Male Kidney TG uptake)

| | # of animals |
|---|---|
| LFD | |
| 16 weeks | 8 |
| 1 year | 8 |
| n3D | |
| 16 weeks | 8 |
| 1 year | 7 (8-1 mouse died=7) |
| WD | |
| 16 weeks | 8 |
| 1 year | 7 (8-1 mouse died=7) |
| | P value |
| 2-way ANOVA | |
| Age | 0.017* |
| Diet | 0.054 |
| Tukey's multiple comparison | |
| 16 weeks | |
| LFD vs ω3FD | 0.422 |
| LFD vs WD | 0.362 |
| ω3FD vs WD | 0.993 |
| 1 year | |
| LFD vs ω3FD | 0.673 |
| LFD vs WD | 0.098 |
| ω3FD vs WD | 0.442 |
| LFD 16 weeks vs 1 year | 0.193 |
| ω3FD 16 weeks vs 1 year | 0.370 |
| WD 16 weeks vs 1 year | 0.045* |

Panel D (Male Quadriceps TG uptake)

| | # of animals |
|---|---|
| LFD | |
| 16 weeks | 8 |
| 1 year | 8 |
| n3D | |

| | |
|---|---|
| 16 weeks | 8 |
| 1 year | 7 (8-1 mouse died=7) |
| WD | |
| 16 weeks | 8 |
| 1 year | 7 (8-1 mouse died=7) |
| | P value |
| 2-way ANOVA | |
| Age | <0.001*** |
| Diet | 0.002** |
| Tukey's multiple comparison | |
| 16 weeks | |
| LFD vs ω3FD | 0.044* |
| LFD vs WD | 0.907 |
| ω3FD vs WD | 0.108 |
| 1 year | |
| LFD vs ω3FD | 0.079 |
| LFD vs WD | 0.920 |
| ω3FD vs WD | 0.040* |
| LFD 16 weeks vs 1 year | 0.032* |
| ω3FD 16 weeks vs 1 year | 0.025* |
| WD 16 weeks vs 1 year | 0.005** |

Panel E (Male gWAT TG uptake)

| | # of animals |
|---|---|
| LFD | |
| 16 weeks | 8 |
| 1 year | 8 |
| n3D | |
| 16 weeks | 8 |
| 1 year | 7 (8-1 mouse died=7) |
| WD | |
| 16 weeks | 8 |
| 1 year | 7 (8-1 mouse died=7) |
| | P value |
| 2-way ANOVA | |
| Age | <0.001*** |
| Diet | <0.001*** |
| Tukey's multiple comparison | |
| 16 weeks | |
| LFD vs ω3FD | 0.328 |
| LFD vs WD | <0.001*** |
| ω3FD vs WD | 0.013* |
| 1 year | |
| LFD vs ω3FD | 0.042* |
| LFD vs WD | 0.003** |
| ω3FD vs WD | 0.586 |
| LFD 16 weeks vs 1 year | 0.002** |
| ω3FD 16 weeks vs 1 year | <0.001*** |
| WD 16 weeks vs 1 year | 0.020* |

Panel F (Male sWAT TG uptake)

| | # of animals |
|---|---|
| LFD | |
| 16 weeks | 8 |
| 1 year | 8 |
| n3D | |
| 16 weeks | 8 |
| 1 year | 7 (8-1 mouse died=7) |
| WD | |
| 16 weeks | 8 |
| 1 year | 7 (8-1 mouse died=7) |
| | P value |
| 2-way ANOVA | |
| Age | <0.001*** |
| Diet | 0.005** |
| Tukey's multiple comparison | |
| 16 weeks | |
| LFD vs ω3FD | 0.215 |
| LFD vs WD | 0.185 |
| ω3FD vs WD | 0.003** |
| 1 year | |
| LFD vs ω3FD | 0.090 |
| LFD vs WD | 0.025* |
| ω3FD vs WD | 0.852 |
| LFD 16 weeks vs 1 year | 0.065 |
| ω3FD 16 weeks vs 1 year | <0.001*** |
| WD 16 weeks vs 1 year | 0.007** |

Panel G (Male BAT TG uptake)

| | # of animals |
|---|---|
| LFD | |
| 16 weeks | 8 |
| 1 year | 8 |
| n3D | |
| 16 weeks | 8 |
| 1 year | 7 (8-1 mouse died=7) |
| WD | |
| 16 weeks | 8 |
| 1 year | 7 (8-1 mouse died=7) |
| | P value |
| 2-way ANOVA | |
| Age | <0.001*** |
| Diet | <0.001*** |
| Tukey's multiple comparison | |
| 16 weeks | |
| LFD vs ω3FD | 0.033* |
| LFD vs WD | 0.074 |
| ω3FD vs WD | <0.001*** |
| 1 year | |
| LFD vs ω3FD | 0.603 |
| LFD vs WD | 0.226 |
| ω3FD vs WD | 0.037* |
| LFD 16 weeks vs 1 year | 0.013* |

| | |
|---|---|
| ω3FD 16 weeks vs 1 year | <0.001*** |
| WD 16 weeks vs 1 year | 0.051 |

**Figure 9 statistics**

Panel A (Female GTT 14 weeks)

| | # of animals |
|---|---|
| LFD | 8 |
| n3D | 8 |
| WD | 8 |
| | P-Value |
| 1-Way ANOVA of AUC | <0.001*** |
| Tukey's multiple comparison | |
| LFD vs ω3FD | 0.007** |
| LFD vs WD | <0.001*** |
| ω3FD vs WD | 0.445 |

Panel B (Female GTT 50 weeks)

| | # of animals |
|---|---|
| LFD | 7 (8-1 mouse died=7) |
| n3D | 6 (8-2 mice died=6) |
| WD | 7 (8-1 mouse died=7) |
| | P-Value |
| 1-Way ANOVA of AUC | 0.296 |
| Tukey's multiple comparison | |
| LFD vs ω3FD | 0.753 |
| LFD vs WD | 0.266 |
| ω3FD vs WD | 0.690 |

Panel C (Female GTT 1.5 years)

| | # of animals |
|---|---|
| LFD | 8 (10-2 mice died=8) |
| n3D | 8 (10-2 mice died=8) |
| WD | 5 (10-5 mice died=5) |
| | P-Value |
| 1-Way ANOVA of AUC | 0.057 |
| Tukey's multiple comparison | |
| LFD vs ω3FD | 0.490 |
| LFD vs WD | 0.046* |
| ω3FD vs WD | 0.281 |

Panel D (Male GTT 14 weeks)

| | # of animals |
|---|---|
| LFD | 8 |
| n3D | 8 |
| WD | 8 |
| | P-Value |
| 1-Way ANOVA of AUC | 0.001** |
| Tukey's multiple comparison | |
| LFD vs ω3FD | 0.623 |
| LFD vs WD | 0.001** |
| ω3FD vs WD | 0.012* |

Panel E (Male GTT 50 weeks)

| | # of animals |
|---|---|
| LFD | 8 |
| n3D | 7 (8-1 mouse died=7) |
| WD | 8 |
| | P-Value |
| 1-Way ANOVA of AUC | 0.139 |
| Tukey's multiple comparison | |
| LFD vs ω3FD | 0.518 |
| LFD vs WD | 0.580 |
| ω3FD vs WD | 0.118 |

Panel F (Male GTT 1.5 years)

| | # of animals |
|---|---|
| LFD | 8 |
| n3D | 7 (8-1 mouse died=7) |
| WD | 5 (8-3 mice died=5) |
| | P-Value |
| 1-Way ANOVA of AUC | 0.038* |
| Tukey's multiple comparison | |
| LFD vs ω3FD | 0.325 |
| LFD vs WD | 0.292 |
| ω3FD vs WD | 0.030* |

Panel G (female insulin 14 weeks)

| | # of animals |
|---|---|
| LFD | 8 |
| n3D | 8 |
| WD | 7 (8-1 sample lost=7) |
| | P value |
| 2-way ANOVA | |
| Time | <0.001*** |
| Diet | 0.004** |
| Tukey's multiple comparison | |
| 0 minutes | |
| LFD vs ω3FD | 0.036* |
| LFD vs WD | 0.999 |
| ω3FD vs WD | 0.050* |
| 30 minutes | |
| LFD vs ω3FD | 0.011* |
| LFD vs WD | 0.778 |
| ω3FD vs WD | 0.002** |
| LFD 0 vs 30 min | 0.026* |
| ω3FD 0 vs 30 min | 0.102 |
| WD 0 vs 30 min | 0.004** |

Panel H (female insulin 50 weeks)

| | # of animals |
|---|---|
| LFD | 7 (8-1 mouse died=7) |
| n3D | 6 (8-2 mice died=6) |
| WD | 7 (8-1 mouse died=7) |
| | P value |
| 2-way ANOVA | |
| Time | <0.001*** |

| | |
|---|---|
| Diet | 0.0016 |
| Tukey’s multiple comparison | |
| 0 minutes | |
| LFD vs ω3FD | 0.071 |
| LFD vs WD | 0.184 |
| ω3FD vs WD | <0.001*** |
| 30 minutes | |
| LFD vs ω3FD | 0.270 |
| LFD vs WD | 0.020* |
| ω3FD vs WD | <0.001*** |
| LFD 0 vs 30 min | 0.017* |
| ω3FD 0 vs 30 min | <0.001*** |
| WD 0 vs 30 min | <0.001*** |

Panel I (female insulin 1.5 years)

| | # of animals |
|---|---|
| LFD | 8 (10-2 mice died=8) |
| n3D | 8 (10-2 mice died=8) |
| WD | 5 (10-5 mice died=5) |
| | P value |
| 2-way ANOVA | |
| Time | <0.001*** |
| Diet | <0.001*** |
| Tukey’s multiple comparison | |
| 0 minutes | |
| LFD vs ω3FD | 0.209 |
| LFD vs WD | 0.019* |
| ω3FD vs WD | <0.001*** |
| 30 minutes | |
| LFD vs ω3FD | 0.520 |
| LFD vs WD | <0.001*** |
| ω3FD vs WD | <0.001*** |
| LFD 0 vs 30 min | 0.008** |
| ω3FD 0 vs 30 min | <0.001*** |
| WD 0 vs 30 min | <0.001*** |

**Figure 10 statistics**

Panel A (Female ITT 15 weeks)

| | # of animals |
|---|---|
| LFD | 8 |
| n3D | 8 |
| WD | 8 |
| | P-Value |
| 1-Way ANOVA of AUC | 0.177 |
| Tukey’s multiple comparison | |
| LFD vs ω3FD | 0.793 |
| LFD vs WD | 0.434 |
| ω3FD vs WD | 0.161 |

Panel B (Female ITT 51 weeks)

| | # of animals |
|---|---|

| | |
|---|---|
| LFD | 7 (8-1 mouse died=7) |
| n3D | 6 (8-2 mice died=6) |
| WD | 7 (8-1 mouse died=7) |
| | P-Value |
| 1-Way ANOVA of AUC | 0.282 |
| Tukey's multiple comparison | |
| LFD vs ω3FD | 0.969 |
| LFD vs WD | 0.291 |
| ω3FD vs WD | 0.373 |

Panel C (Female ITT 1.5 years)

| | # of animals |
|---|---|
| LFD | 5 (10-2 mice died-3 mice didn't finish=5) |
| n3D | 2 (10-2 mice died-6 mice didn't finish=2) |
| WD | 5 (10-5 mice died=5) |
| | P-Value |
| 1-Way ANOVA of AUC | 0.627 |
| Tukey's multiple comparison | |
| LFD vs ω3FD | 0.967 |
| LFD vs WD | 0.714 |
| ω3FD vs WD | 0.684 |

Panel D (Male ITT 15 weeks)

| | # of animals |
|---|---|
| LFD | 8 |
| n3D | 8 |
| WD | 8 |
| | P-Value |
| 1-Way ANOVA of AUC | <0.001*** |
| Tukey's multiple comparison | |
| LFD vs ω3FD | 0.040* |
| LFD vs WD | 0.186 |
| ω3FD vs WD | <0.001*** |

Panel E (Male ITT 51 weeks)

| | # of animals |
|---|---|
| LFD | 8 |
| n3D | 7 (8-1 mouse died=7) |
| WD | 8 |
| | P-Value |
| 1-Way ANOVA of AUC | 0.025* |
| Tukey's multiple comparison | |
| LFD vs ω3FD | 0.283 |
| LFD vs WD | 0.324 |
| ω3FD vs WD | 0.019* |

Panel F (Male ITT 1.5 years)

| # of animal | |
|---|---|
| LFD | 8 |
| n3D | 6 (8-1 mouse died-1 outlier value=6) |
| WD | 5 (8-3 mice died=5) |
| | P-Value |
| 1-Way ANOVA of AUC | 0.272 |
| Tukey's multiple comparison | |

| | |
|---|---|
| LFD vs ω3FD | 0.994 |
| LFD vs WD | 0.320 |
| ω3FD vs WD | 0.318 |

**Supplementary Figure 1 statistics**

Panel A (Female Lean Mass)

| # of animals | # of animals |
|---|---|
| LFD | |
| 8 weeks (pre-diet) | 26 |
| 16 weeks | 26 |
| 8 months | 16 (18-2 mice died=16) |
| 1 year | 15 (18-3 mice died=15) |
| 1.5 year | 8 (10-2 mice died=8) |
| 2 years | 6 (10-4 mice died=6) |
| n3D | |
| 8 weeks (pre-diet) | 25 (26-1 outlier value) |
| 16 weeks | 26 |
| 8 months | 15 (18-3 mice died=15) |
| 1 year | 14 (18-4 mice died=14) |
| 1.5 year | 7 (10-3 mice died=7) |
| 2 years | 4 (10-6 mice died=4) |
| WD | |
| 8 weeks (pre-diet) | 26 |
| 16 weeks | 26 |
| 8 months | 17 (18-1 mouse died=17 |
| 1 year | 15 (18-3 mice died=15) |
| 1.5 year | 5 (10-5 mice died=5) |
| 2 years | 2 (10-8 mice died=2) |

Panel B (Male Lean Mass)

| | # of animals |
|---|---|
| LFD | |
| 8 weeks (pre-diet) | 24 |
| 16 weeks | 24 |
| 8 months | 16 |
| 1 year | 16 |
| 1.5 year | 8 |
| 2 years | 6 (8-2 mice died=6) |
| n3D | |
| 8 weeks (pre-diet) | 24 |
| 16 weeks | 24 |
| 8 months | 14 (16-1 mouse died-1 mouse not measured=14) |
| 1 year | 15 (16-1 mouse died) |
| 1.5 year | 7 (8-1 mouse died=7) |
| 2 years | 5 (8-3 mice died=5) |
| WD | |
| 8 weeks (pre-diet) | 24 |
| 16 weeks | 24 |
| 8 months | 16 |
| 1 year | 14 (16-2 mice not measured=14) |

| | |
|---|---|
| 1.5 year | 5 (8-3 mice died=5) |
| 2 years | 3 (8-5 mice died=3) |

Panel C (Female Fat Mass)

| | # of animals |
|---|---|
| LFD | |
| 8 weeks (pre-diet) | 26 |
| 16 weeks | 26 |
| 8 months | 16 (18-2 mice died=16) |
| 1 year | 15 (18-3 mice died=15) |
| 1.5 year | 8 (10-2 mice died=8) |
| 2 years | 6 (10-4 mice died=6) |
| n3D | |
| 8 weeks (pre-diet) | 26 |
| 16 weeks | 26 |
| 8 months | 15 (18-3 mice died=15) |
| 1 year | 14 (18-4 mice died=14) |
| 1.5 year | 7 (10-3 mice died=7) |
| 2 years | 4 (10-6 mice died=4) |
| WD | |
| 8 weeks (pre-diet) | 26 |
| 16 weeks | 24 (26-2 outlier values) |
| 8 months | 17 (18-1 mouse died=17) |
| 1 year | 15 (18-3 mice died=15) |
| 1.5 year | 5 (10-5 mice died=5) |
| 2 years | 2 (10-8 mice died=2) |

Panel D (Male Fat Mass)

| | # of animals |
|---|---|
| LFD | |
| 8 weeks (pre-diet) | 24 |
| 16 weeks | 23 (24-1 outlier value=23) |
| 8 months | 16 |
| 1 year | 16 |
| 1.5 year | 8 |
| 2 years | 6 (8-2 mice died=6) |
| n3D | |
| 8 weeks (pre-diet) | 24 |
| 16 weeks | 24 |
| 8 months | 14 (16-1 mouse died-1 mouse not measured=14) |
| 1 year | 15 (16-1 mouse died) |
| 1.5 year | 7 (8-1 mouse died=7) |
| 2 years | 5 (8-3 mice died=5) |
| WD | |
| 8 weeks (pre-diet) | 24 |
| 16 weeks | 24 |
| 8 months | 16 |
| 1 year | 14 (16-2 mice not measured) |
| 1.5 year | 5 (8-3 mice died=5) |
| 2 years | 3 (8-5 mice died=3) |

**Supplementary Figure 2 statistics**

Panel A (Plasma TG females)

| | # of animals |
|---|---|
| LFD | |
| 8 weeks (pre-diet) | 10 |
| 16 weeks | 10 |
| 8 months | 9 (10-1 mouse died=9) |
| 1 year | 8 (10-2 mice died=8) |
| 1.5 year | 8 (10-2 mice died=8) |
| 2 years | 6 (10-4 mice died=6) |
| n3D | |
| 8 weeks (pre-diet) | 10 |
| 16 weeks | 10 |
| 8 months | 9 (10-1 mouse died=9) |
| 1 year | 8 (10-2 mice died=8) |
| 1.5 year | 8 (10-2 mice died=8) |
| 2 years | 4 (10-6 mice died=4) |
| WD | |
| 8 weeks (pre-diet) | 10 |
| 16 weeks | 10 |
| 8 months | 10 |
| 1 year | 7 (10-2 mice died-1 outlier value=7) |
| 1.5 year | 5 (10-5 mice died=5) |
| 2 years | 2 (10-8 mice died=2) |

Panel B (Plasma TG males)

| | # of animals |
|---|---|
| LFD | |
| 8 weeks (pre-diet) | 8 |
| 16 weeks | 8 |
| 8 months | 8 |
| 1 year | 8 |
| 1.5 year | 8 |
| 2 years | 6 (8-2 mice died=6) |
| n3D | |
| 8 weeks (pre-diet) | 8 |
| 16 weeks | 8 |
| 8 months | 7 (8-1 sample lost=7) |
| 1 year | 8 |
| 1.5 year | 7 (8-1 mouse died=7) |
| 2 years | 5 (8-3 mice died=5) |
| WD | |
| 8 weeks (pre-diet) | 8 |
| 16 weeks | 8 |
| 8 months | 8 |
| 1 year | 7 (8-1 sample lost=7) |
| 1.5 year | 5 (8-3 mice died=5) |
| 2 years | 3 (8-5 mice died=3) |

**Supplementary Figure 3 statistics**

Panel A (Female FTT 2 years)

| | # of animals |
|---|---|
| LFD | 6 (10-4 mice died=6) |
| n3D | 4 (10-6 mice died=4) |
| WD | 2 (10-8 mice died=2) |

Panel B (Male FTT 2 years)

| | # of animals |
|---|---|
| LFD | 6 (8-2 mice died=6) |
| n3D | 5 (8-3 mice died=5) |
| WD | 3 (8-5 mice died=3) |

**Supplementary Figure 4 statistics**

Panel A (Female 2-year chylomicron clearance)

| | # of animals |
|---|---|
| LFD | 6 (10-4 mice died=6) |
| n3D | 3 (10-7 mice died=3) |
| WD | 0 (10-10 mice died=0) |

Panel B (Male 2-year chylomicron clearance)

| | # of animals |
|---|---|
| LFD | 5 (8-3 mice died=5) |
| n3D | 5 (8-3 mice died=5) |
| WD | 3 (8-5 mice died=3) |

**Supplementary Figure 5 statistics**

Panel A (Female Heart TG uptake)

| | # of animals |
|---|---|
| LFD | |
| 16 weeks | 8 |
| 1 year | 7 (8-1 mouse died=7) |
| n3D | |
| 16 weeks | 8 |
| 1 year | 6 (8-2 mice died=6) |
| WD | |
| 16 weeks | 8 |
| 1 year | 7 (8-1 mouse died=7) |
| | P value |
| 2-way ANOVA | |
| Age | 0.129 |
| Diet | <0.001*** |
| Tukey’s multiple comparison | |
| 16 weeks | |
| LFD vs n3D | <0.001*** |
| LFD vs WD | 0.390 |
| n3D vs WD | <0.001*** |
| 1 year | |
| LFD vs n3D | 0.005** |
| LFD vs WD | 0.993 |

| | |
|---|---|
| n3D vs WD | 0.004** |
| LFD | |
| 16 weeks vs. 1 year | 0.059 |
| n3D | |
| 16 weeks vs. 1 year | 0.826 |
| WD | |
| 16 weeks vs. 1 year | 0.586 |

Panel B (Female Liver TG uptake)

| | # of animals |
|---|---|
| LFD | |
| 16 weeks | 8 |
| 1 year | 7 (8-1 mouse died=7) |
| n3D | |
| 16 weeks | 8 |
| 1 year | 6 (8-2 mice died=6) |
| WD | |
| 16 weeks | 8 |
| 1 year | 7 (8-1 mouse died=7) |
| | P value |
| 2-way ANOVA | |
| Age | 0.004** |
| Diet | 0.079 |
| Tukey's multiple comparison | |
| 16 weeks | |
| LFD vs n3D | 0.082 |
| LFD vs WD | 0.026* |
| n3D vs WD | 0.869 |
| 1 year | |
| LFD vs n3D | 0.536 |
| LFD vs WD | 0.896 |
| n3D vs WD | 0.301 |
| LFD | |
| 16 weeks vs. 1 year | 0.680 |
| n3D | |
| 16 weeks vs. 1 year | 0.188 |
| WD | |
| 16 weeks vs. 1 year | 0.001** |

Panel C (Female Kidney TG uptake)

| | # of animals |
|---|---|
| LFD | |
| 16 weeks | 8 |
| 1 year | 7 (8-1 mouse died=7) |
| n3D | |
| 16 weeks | 7 (8-1 outlier value=7) |
| 1 year | 6 (8-2 mice died=6) |
| WD | |
| 16 weeks | 8 |
| 1 year | 7 (8-1 mouse died=7) |
| | P value |
| 2-way ANOVA | |
| Age | 0.562 |
| Diet | <0.001*** |

| | |
|---|---|
| Tukey's multiple comparison | |
| 16 weeks | |
| LFD vs n3D | 0.003** |
| LFD vs WD | 0.794 |
| n3D vs WD | 0.016* |
| 1 year | |
| LFD vs n3D | 0.769 |
| LFD vs WD | 0.043 |
| n3D vs WD | 0.010* |
| LFD | |
| 16 weeks vs. 1 year | 0.105 |
| n3D | |
| 16 weeks vs. 1 year | 0.293 |
| WD | |
| 16 weeks vs. 1 year | 0.129 |

Panel D (Female Quadriceps TG uptake)

| | # of animals |
|---|---|
| LFD | |
| 16 weeks | 8 |
| 1 year | 7 (8-1 mouse died=7) |
| n3D | |
| 16 weeks | 7 (8-1 outlier value=7) |
| 1 year | 6 (8-2 mice died=6) |
| WD | |
| 16 weeks | 8 |
| 1 year | 7 (8-1 mouse died=7) |
| | P value |
| 2-way ANOVA | |
| Age | 0.027* |
| Diet | 0.870 |
| Tukey's multiple comparison | |
| 16 weeks | |
| LFD vs n3D | 0.994 |
| LFD vs WD | 0.446 |
| n3D vs WD | 0.530 |
| 1 year | |
| LFD vs n3D | 0.933 |
| LFD vs WD | 0.902 |
| n3D vs WD | 0.723 |
| LFD | |
| 16 weeks vs. 1 year | 0.078 |
| n3D | |
| 16 weeks vs. 1 year | 0.059 |
| WD | |
| 16 weeks vs. 1 year | 0.858 |

Panel E (Female gWAT TG uptake)

| | # of animals |
|---|---|
| LFD | |
| 16 weeks | 8 |
| 1 year | 7 (8-1 mouse died=7) |
| n3D | |
| 16 weeks | 8 |

| | |
|---|---|
| 1 year | 6 (8-2 mice died=6) |
| WD | |
| 16 weeks | 8 |
| 1 year | 7 (8-1 mouse died=7) |
| | P value |
| 2-way ANOVA | |
| Age | 0.006** |
| Diet | 0.325 |
| Tukey's multiple comparison | |
| 16 weeks | |
| LFD vs n3D | 0.139 |
| LFD vs WD | 0.961 |
| n3D vs WD | 0.081 |
| 1 year | |
| LFD vs n3D | 0.796 |
| LFD vs WD | 0.750 |
| n3D vs WD | 0.999 |
| LFD | |
| 16 weeks vs. 1 year | 0.009** |
| n3D | |
| 16 weeks vs. 1 year | 0.880 |
| WD | |
| 16 weeks vs. 1 year | 0.031* |

Panel F (Female BAT TG uptake)

| | # of animals |
|---|---|
| LFD | |
| 16 weeks | 8 |
| 1 year | 7 (8-1 mouse died=7) |
| n3D | |
| 16 weeks | 8 |
| 1 year | 6 (8-2 mice died=6) |
| WD | |
| 16 weeks | 8 |
| 1 year | 7 (8-1 mouse died=7) |
| | P value |
| 2-way ANOVA | |
| Age | 0.014* |
| Diet | 0.102 |
| Tukey's multiple comparison | |
| 16 weeks | |
| LFD vs n3D | 0.860 |
| LFD vs WD | 0.347 |
| n3D vs WD | 0.653 |
| 1 year | |
| LFD vs n3D | 0.408 |
| LFD vs WD | 0.705 |
| n3D vs WD | 0.112 |
| LFD | |
| 16 weeks vs. 1 year | 0.488 |
| n3D | |
| 16 weeks vs. 1 year | 0.017* |
| WD | |

| | |
|---|---|
| 16 weeks vs. 1 year | 0.225 |

Panel G (Male Heart TG uptake)

| | # of animals |
|---|---|
| LFD | |
| 16 weeks | 8 |
| 1 year | 8 |
| n3D | |
| 16 weeks | 7 (8-1 outlier value=7) |
| 1 year | 7 (8-1 mouse died=7) |
| WD | |
| 16 weeks | 8 |
| 1 year | 6 (8-2 outlier values=6) |
| | P value |
| 2-way ANOVA | |
| Age | 0.060 |
| Diet | <0.001*** |
| Tukey’s multiple comparison | |
| 16 weeks | |
| LFD vs n3D | <0.001*** |
| LFD vs WD | 0.984 |
| n3D vs WD | <0.001*** |
| 1 year | |
| LFD vs n3D | <0.001*** |
| LFD vs WD | 0.895 |
| n3D vs WD | <0.001*** |
| LFD | |
| 16 weeks vs. 1 year | 0.703 |
| n3D | |
| 16 weeks vs. 1 year | 0.028* |
| WD | |
| 16 weeks vs. 1 year | 0.522 |

Panel H (Male Liver TG uptake)

| | # of animals |
|---|---|
| LFD | |
| 16 weeks | 8 |
| 1 year | 8 |
| n3D | |
| 16 weeks | 8 |
| 1 year | 6 (8-1 mouse died-1 outlier value=6) |
| WD | |
| 16 weeks | 8 |
| 1 year | 8 |
| | P value |
| 2-way ANOVA | |
| Age | <0.001*** |
| Diet | 0.002** |
| Tukey’s multiple comparison | |
| 16 weeks | |
| LFD vs n3D | 0.542 |
| LFD vs WD | 0.984 |
| n3D vs WD | 0.441 |
| 1 year | |

| | |
|---|---|
| LFD vs n3D | 0.068 |
| LFD vs WD | 0.122 |
| n3D vs WD | <0.001*** |
| LFD | |
| 16 weeks vs. 1 year | <0.001*** |
| n3D | |
| 16 weeks vs. 1 year | 0.020* |
| WD | |
| 16 weeks vs. 1 year | <0.001*** |

Panel I (Male Kidney TG uptake)

| | # of animals |
|---|---|
| LFD | |
| 16 weeks | 8 |
| 1 year | 8 |
| n3D | |
| 16 weeks | 8 |
| 1 year | 7 (8-1 mouse died=7) |
| WD | |
| 16 weeks | 8 |
| 1 year | 8 |
| | P value |
| 2-way ANOVA | |
| Age | 0.086 |
| Diet | 0.109 |
| Tukey's multiple comparison | |
| 16 weeks | |
| LFD vs n3D | 0.982 |
| LFD vs WD | 0.608 |
| n3D vs WD | 0.495 |
| 1 year | |
| LFD vs n3D | 0.671 |
| LFD vs WD | 0.563 |
| n3D vs WD | 0.166 |
| LFD | |
| 16 weeks vs. 1 year | 0.419 |
| n3D | |
| 16 weeks vs. 1 year | 0.150 |
| WD | |
| 16 weeks vs. 1 year | 0.461 |

Panel J (Male Quadriceps TG uptake)

| | # of animals |
|---|---|
| LFD | |
| 16 weeks | 8 |
| 1 year | 8 |
| n3D | |
| 16 weeks | 8 |
| 1 year | 7 (8-1 mouse died=7) |
| WD | |
| 16 weeks | 8 |
| 1 year | 8 |
| | P value |
| 2-way ANOVA | |

| | |
|---|---|
| Age | 0.032* |
| Diet | <0.001*** |
| Tukey's multiple comparison | |
| 16 weeks | |
| LFD vs n3D | 0.153 |
| LFD vs WD | 0.999 |
| n3D vs WD | 0.157 |
| 1 year | |
| LFD vs n3D | 0.025* |
| LFD vs WD | 0.257 |
| n3D vs WD | <0.001*** |
| LFD | |
| 16 weeks vs. 1 year | 0.291 |
| n3D | |
| 16 weeks vs. 1 year | 0.893 |
| WD | |
| 16 weeks vs. 1 year | 0.010* |

Panel K (Male gWAT TG uptake)

| | # of animals |
|---|---|
| LFD | |
| 16 weeks | 8 |
| 1 year | 8 |
| n3D | |
| 16 weeks | 8 |
| 1 year | 7 (8-1 mouse died=7) |
| WD | |
| 16 weeks | 8 |
| 1 year | 8 |
| | P value |
| 2-way ANOVA | |
| Age | <0.001*** |
| Diet | 0.455 |
| Tukey's multiple comparison | |
| 16 weeks | |
| LFD vs n3D | 0.527 |
| LFD vs WD | 0.431 |
| n3D vs WD | 0.062 |
| 1 year | |
| LFD vs n3D | 0.778 |
| LFD vs WD | 0.180 |
| n3D vs WD | 0.539 |
| LFD | |
| 16 weeks vs. 1 year | <0.001*** |
| n3D | |
| 16 weeks vs. 1 year | <0.001*** |
| WD | |
| 16 weeks vs. 1 year | 0.268 |

Panel L (Male BAT TG uptake)

| | # of animals |
|---|---|
| LFD | |
| 16 weeks | 8 |
| 1 year | 8 |

| | |
|---|---|
| n3D | |
| 16 weeks | 8 |
| 1 year | 7 (8-1 mouse died=7) |
| WD | |
| 16 weeks | 8 |
| 1 year | 8 |
| | P value |
| 2-way ANOVA | |
| Age | 0.057 |
| Diet | <0.001*** |
| Tukey's multiple comparison | |
| 16 weeks | |
| LFD vs n3D | 0.561 |
| LFD vs WD | 0.302 |
| n3D vs WD | 0.040* |
| 1 year | |
| LFD vs n3D | 0.001** |
| LFD vs WD | 0.603 |
| n3D vs WD | <0.001*** |
| LFD | |
| 16 weeks vs. 1 year | 0.986 |
| n3D | |
| 16 weeks vs. 1 year | 0.007** |
| WD | |
| 16 weeks vs. 1 year | 0.608 |

**Supplementary Figure 6 statistics**

Panel A (Female Heart TG uptake)

| | # of animals |
|---|---|
| LFD | |
| 16 weeks | 8 |
| 1 year | 6 (8-1 mouse died-one outlier value=6) |
| 2 years | 6 (10-4 mice died=6) |
| n3D | |
| 16 weeks | 7 (8-1 outlier value=7) |
| 1 year | 6 (8-2 mice died=6) |
| 2 years | 3 (10-7 mice died=3) |
| WD | |
| 16 weeks | 8 |
| 1 year | 7 (8-1 mouse died=7) |
| 2 years | 0 (10-10 mice died=0) |
| | P value |
| 2-way ANOVA | |
| Age | <0.001*** |
| Diet | <0.001*** |
| Tukey's multiple comparison | |
| 16 weeks | |
| LFD vs n3D | <0.001*** |
| LFD vs WD | 0.512 |
| n3D vs WD | <0.001*** |
| 1 year | |

| | |
|---|---|
| LFD vs n3D | 0.043* |
| LFD vs WD | 0.820 |
| n3D vs WD | 0.007** |
| 2 years | |
| LFD vs n3D | 0.009** |
| LFD | |
| 16 weeks vs 1 year | 0.959 |
| 16 weeks vs 2 years | 0.983 |
| 1 year vs 2 years | 0.995 |
| n3D | |
| 16 weeks vs 1 year | <0.001*** |
| 16 weeks vs 2 years | 0.006** |
| 1 year vs 2 years | 0.604 |
| WD | |
| 16 weeks vs. 1 year | 0.332 |

Panel B (Female Liver TG uptake)

| | # of animals |
|---|---|
| LFD | |
| 16 weeks | 8 |
| 1 year | 7 (8-1 mouse died=7) |
| 2 years | 6 (10-4 mice died=6) |
| n3D | |
| 16 weeks | 8 |
| 1 year | 6 (8-2 mice died=6) |
| 2 years | 3 (10-7 mice died=3) |
| WD | |
| 16 weeks | 8 |
| 1 year | 7 (8-1 mouse died=7) |
| 2 years | 0 (10-10 mice died=0) |
| | P value |
| 2-way ANOVA | |
| Age | 0.001** |
| Diet | <0.001*** |
| Tukey's multiple comparison | |
| 16 weeks | |
| LFD vs n3D | <0.001*** |
| LFD vs WD | <0.001*** |
| n3D vs WD | 0.952 |
| 1 year | |
| LFD vs n3D | 0.045* |
| LFD vs WD | 0.028* |
| n3D vs WD | 0.996 |
| 2 years | |
| LFD vs n3D | 0.003** |
| LFD | |
| 16 weeks vs 1 year | 0.008** |
| 16 weeks vs 2 years | 0.093 |
| 1 year vs 2 years | 0.674 |
| n3D | |
| 16 weeks vs 1 year | 0.195 |
| 16 weeks vs 2 years | 0.067 |
| 1 year vs 2 years | 0.674 |

| | |
|---|---|
| WD | |
| 16 weeks vs. 1 year | 0.238 |

Panel C (Female Kidney TG uptake)

| | # of animals |
|---|---|
| LFD | |
| 16 weeks | 8 |
| 1 year | 7 (8-1 mouse died=7) |
| 2 years | 6 (10-4 mice died=6) |
| n3D | |
| 16 weeks | 8 |
| 1 year | 6 (8-2 mice died=6) |
| 2 years | 3 (10-7 mice died=3) |
| WD | |
| 16 weeks | 8 |
| 1 year | 7 (8-1 mouse died=7) |
| 2 years | 0 (10-10 mice died=0) |
| | P value |
| 2-way ANOVA | |
| Age | <0.001*** |
| Diet | 0.480 |
| Tukey's multiple comparison | |
| 16 weeks | |
| LFD vs n3D | 0.136 |
| LFD vs WD | 0.903 |
| n3D vs WD | 0.290 |
| 1 year | |
| LFD vs n3D | 0.145 |
| LFD vs WD | 0.032* |
| n3D vs WD | 0.828 |
| 2 years | |
| LFD vs n3D | 0.262 |
| LFD | |
| 16 weeks vs 1 year | 0.992 |
| 16 weeks vs 2 years | 0.574 |
| 1 year vs 2 years | 0.522 |
| n3D | |
| 16 weeks vs 1 year | <0.001*** |
| 16 weeks vs 2 years | 0.067 |
| 1 year vs 2 years | 0.713 |
| WD | |
| 16 weeks vs. 1 year | 0.006** |

Panel D (Female Quadriceps TG uptake)

| | # of animals |
|---|---|
| LFD | |
| 16 weeks | 8 |
| 1 year | 7 (8-1 mouse died=7) |
| 2 years | 6 (10-4 mice died=6) |
| n3D | |
| 16 weeks | 7 (8-1 outlier value=7) |
| 1 year | 6 (8-2 mice died=6) |
| 2 years | 3 (10-7 mice died=3) |
| WD | |

| | |
|---|---|
| 16 weeks | 8 |
| 1 year | 7 (8-1 mouse died=7) |
| 2 years | 0 (10-10 mice died=0) |
| | P value |
| 2-way ANOVA | |
| Age | <0.001*** |
| Diet | 0.738 |
| Tukey's multiple comparison | |
| 16 weeks | |
| LFD vs n3D | 0.302 |
| LFD vs WD | 0.357 |
| n3D vs WD | 0.986 |
| 1 year | |
| LFD vs n3D | 0.995 |
| LFD vs WD | 0.795 |
| n3D vs WD | 0.859 |
| 2 years | |
| LFD vs n3D | 0.297 |
| LFD | |
| 16 weeks vs 1 year | 0.302 |
| 16 weeks vs 2 years | <0.001*** |
| 1 year vs 2 years | 0.003** |
| n3D | |
| 16 weeks vs 1 year | 0.995 |
| 16 weeks vs 2 years | 0.402 |
| 1 year vs 2 years | 0.379 |
| WD | |
| 16 weeks vs. 1 year | 0.867 |

Panel E (Female gWAT TG uptake)

| | # of animals |
|---|---|
| LFD | |
| 16 weeks | 8 |
| 1 year | 7 (8-1 mouse died=7) |
| 2 years | 6 (10-4 mice died=6) |
| n3D | |
| 16 weeks | 8 |
| 1 year | 6 (8-2 mice died=6) |
| 2 years | 3 (10-7 mice died=3) |
| WD | |
| 16 weeks | 8 |
| 1 year | 7 (8-1 mouse died=7) |
| 2 years | 0 (10-10 mice died=0) |
| | P value |
| 2-way ANOVA | |
| Age | 0.007** |
| Diet | 0.379 |
| Tukey's multiple comparison | |
| 16 weeks | |
| LFD vs n3D | 0.333 |
| LFD vs WD | 0.473 |
| n3D vs WD | 0.032* |
| 1 year | |

| | |
|---|---|
| LFD vs n3D | 0.102 |
| LFD vs WD | 0.059 |
| n3D vs WD | 0.985 |
| 2 years | |
| LFD vs n3D | 0.958 |
| LFD | |
| 16 weeks vs 1 year | 0.949 |
| 16 weeks vs 2 years | 0.624 |
| 1 year vs 2 years | 0.812 |
| n3D | |
| 16 weeks vs 1 year | 0.001** |
| 16 weeks vs 2 years | 0.297 |
| 1 year vs 2 years | 0.328 |
| WD | |
| 16 weeks vs. 1 year | 0.255 |

Panel F (Female sWAT TG uptake)

| | # of animals |
|---|---|
| LFD | |
| 16 weeks | 8 |
| 1 year | 7 (8-1 mouse died=7) |
| 2 years | 6 (10-4 mice died=6) |
| n3D | |
| 16 weeks | 8 |
| 1 year | 6 (8-2 mice died=6) |
| 2 years | 3 (10-7 mice died=3) |
| WD | |
| 16 weeks | 8 |
| 1 year | 7 (8-1 mouse died=7) |
| 2 years | 0 (10-10 mice died=0) |
| | P value |
| 2-way ANOVA | |
| Age | 0.002** |
| Diet | 0.116 |
| Tukey's multiple comparison | |
| 16 weeks | |
| LFD vs n3D | 0.946 |
| LFD vs WD | 0.483 |
| n3D vs WD | 0.311 |
| 1 year | |
| LFD vs n3D | 0.063 |
| LFD vs WD | 0.040* |
| n3D vs WD | 0.995 |
| 2 years | |
| LFD vs n3D | 0.989 |
| LFD | |
| 16 weeks vs 1 year | 0.884 |
| 16 weeks vs 2 years | 0.902 |
| 1 year vs 2 years | 0.668 |
| n3D | |
| 16 weeks vs 1 year | 0.008** |
| 16 weeks vs 2 years | 0.963 |
| 1 year vs 2 years | 0.029* |

| | |
|---|---|
| WD | |
| 16 weeks vs. 1 year | 0.137 |

Panel G (Female BAT TG uptake)

| | # of animals |
|---|---|
| LFD | |
| 16 weeks | 8 |
| 1 year | 7 (8-1 mouse died=7) |
| 2 years | 6 (10-4 mice died=6) |
| n3D | |
| 16 weeks | 8 |
| 1 year | 6 (8-2 mice died=6) |
| 2 years | 3 (10-7 mice died=3) |
| WD | |
| 16 weeks | 8 |
| 1 year | 7 (8-1 mouse died=7) |
| 2 years | 0 (10-10 mice died=0) |
| | P value |
| 2-way ANOVA | |
| Age | <0.001*** |
| Diet | 0.183 |
| Tukey's multiple comparison | |
| 16 weeks | |
| LFD vs n3D | 0.507 |
| LFD vs WD | 0.779 |
| n3D vs WD | 0.897 |
| 1 year | |
| LFD vs n3D | 0.950 |
| LFD vs WD | 0.069 |
| n3D vs WD | 0.154 |
| 2 years | |
| LFD vs n3D | 0.292 |
| LFD | |
| 16 weeks vs 1 year | 0.863 |
| 16 weeks vs 2 years | 0.382 |
| 1 year vs 2 years | 0.187 |
| n3D | |
| 16 weeks vs 1 year | 0.972 |
| 16 weeks vs 2 years | 0.003** |
| 1 year vs 2 years | 0.007** |
| WD | |
| 16 weeks vs. 1 year | 0.078 |

**Supplementary Figure statistics**

Panel A (Male Heart TG uptake)

| | # of animals |
|---|---|
| LFD | |
| 16 weeks | 7 (8-1 outlier value=7) |
| 1 year | 8 |
| 2 years | 5 (8-3 mice died=5) |
| n3D | |

| | |
|---|---|
| 16 weeks | 8 |
| 1 year | 7 (8-1 mouse died=7) |
| 2 years | 5 (8-3 mice died=5) |
| WD | |
| 16 weeks | 8 |
| 1 year | 7 (8-1 mouse died=7) |
| 2 years | 3 (8-5 mice died=3) |
| | P value |
| 2-way ANOVA | |
| Age | 0.001** |
| Diet | <0.001*** |
| Tukey's multiple comparison | |
| 16 weeks | |
| LFD vs n3D | <0.001*** |
| LFD vs WD | 0.826 |
| n3D vs WD | <0.001*** |
| 1 year | |
| LFD vs n3D | <0.001*** |
| LFD vs WD | 0.979 |
| n3D vs WD | <0.001*** |
| 2 years | |
| LFD vs n3D | 0.015 |
| LFD vs WD | 0.434 |
| n3D vs WD | 0.422 |
| LFD | |
| 16 weeks vs 1 year | 0.943 |
| 16 weeks vs 2 years | 0.113 |
| 1 year vs 2 years | 0.180 |
| n3D | |
| 16 weeks vs 1 year | 0.994 |
| 16 weeks vs 2 years | <0.001*** |
| 1 year vs 2 years | <0.001*** |
| WD | |
| 16 weeks vs. 1 year | 0.509 |
| 16 weeks vs 2 years | 0.657 |
| 1 year vs 2 years | 0.998 |

Panel B (Male Liver TG uptake)

| | # of animals |
|---|---|
| LFD | |
| 16 weeks | 8 |
| 1 year | 8 |
| 2 years | 5 (8-3 mice died=5) |
| n3D | |
| 16 weeks | 8 |
| 1 year | 6 (8-1 mouse died-1 outlier value=6) |
| 2 years | 5 (8-3 mice died=5) |
| WD | |
| 16 weeks | 8 |
| 1 year | 7 (8-1 mouse died=7) |
| 2 years | 3 (8-5 mice died=3) |
| | P value |
| 2-way ANOVA | |

| | |
|---|---|
| Age | <0.001*** |
| Diet | <0.001*** |
| Tukey's multiple comparison | |
| 16 weeks | |
| LFD vs n3D | 0.008** |
| LFD vs WD | 0.039* |
| n3D vs WD | 0.824 |
| 1 year | |
| LFD vs n3D | 0.007** |
| LFD vs WD | <0.001*** |
| n3D vs WD | 0.096 |
| 2 years | |
| LFD vs n3D | 0.289 |
| LFD vs WD | 0.557 |
| n3D vs WD | 0.956 |
| LFD | |
| 16 weeks vs 1 year | 0.973 |
| 16 weeks vs 2 years | <0.001*** |
| 1 year vs 2 years | <0.001*** |
| n3D | |
| 16 weeks vs 1 year | 0.874 |
| 16 weeks vs 2 years | 0.003** |
| 1 year vs 2 years | 0.016* |
| WD | |
| 16 weeks vs. 1 year | 0.004** |
| 16 weeks vs 2 years | 0.009** |
| 1 year vs 2 years | 0.860 |

Panel C (Male Kidney TG uptake)

| | # of animals |
|---|---|
| LFD | |
| 16 weeks | 8 |
| 1 year | 8 |
| 2 years | 5 (8-3 mice died=5) |
| n3D | |
| 16 weeks | 8 |
| 1 year | 7 (8-1 mouse died=7) |
| 2 years | 5 (8-3 mice died=5) |
| WD | |
| 16 weeks | 8 |
| 1 year | 7 (8-1 mouse died=7) |
| 2 years | 3 (8-5 mice died=3) |
| | P value |
| 2-way ANOVA | |
| Age | 0.052 |
| Diet | 0.002** |
| Tukey's multiple comparison | |
| 16 weeks | |
| LFD vs n3D | 0.434 |
| LFD vs WD | 0.374 |
| n3D vs WD | 0.993 |
| 1 year | |
| LFD vs n3D | 0.683 |

| | |
|---|---|
| LFD vs WD | 0.103 |
| n3D vs WD | 0.453 |
| 2 years | |
| LFD vs n3D | 0.273 |
| LFD vs WD | 0.023* |
| n3D vs WD | 0.354 |
| LFD | |
| 16 weeks vs 1 year | 0.403 |
| 16 weeks vs 2 years | 0.762 |
| 1 year vs 2 years | 0.166 |
| n3D | |
| 16 weeks vs 1 year | 0.650 |
| 16 weeks vs 2 years | 0.998 |
| 1 year vs 2 years | 0.674 |
| WD | |
| 16 weeks vs. 1 year | 0.115 |
| 16 weeks vs 2 years | 0.369 |
| 1 year vs 2 years | 0.981 |

Panel D (Male Quadriceps TG uptake)

| | # of animals |
|---|---|
| LFD | |
| 16 weeks | 8 |
| 1 year | 8 |
| 2 years | 5 (8-3 mice died=5) |
| n3D | |
| 16 weeks | 8 |
| 1 year | 7 (8-1 mouse died=7) |
| 2 years | 5 (8-3 mice died=5) |
| WD | |
| 16 weeks | 8 |
| 1 year | 7 (8-1 mouse died=7) |
| 2 years | 3 (8-5 mice died=3) |
| | P value |
| 2-way ANOVA | |
| Age | <0.001*** |
| Diet | 0.003** |
| Tukey's multiple comparison | |
| 16 weeks | |
| LFD vs n3D | 0.028* |
| LFD vs WD | 0.895 |
| n3D vs WD | 0.079 |
| 1 year | |
| LFD vs n3D | 0.056 |
| LFD vs WD | 0.909 |
| n3D vs WD | 0.025* |
| 2 years | |
| LFD vs n3D | 0.954 |
| LFD vs WD | 0.570 |
| n3D vs WD | 0.419 |
| LFD | |
| 16 weeks vs 1 year | 0.056 |
| 16 weeks vs 2 years | 0.714 |

| | |
|---|---|
| 1 year vs 2 years | 0.017* |
| n3D | |
| 16 weeks vs 1 year | 0.043* |
| 16 weeks vs 2 years | 0.446 |
| 1 year vs 2 years | 0.577 |
| WD | |
| 16 weeks vs. 1 year | 0.008** |
| 16 weeks vs 2 years | 0.725 |
| 1 year vs 2 years | 0.256 |

Panel E (Male gWAT TG uptake)

| | # of animals |
|---|---|
| LFD | |
| 16 weeks | 8 |
| 1 year | 8 |
| 2 years | 5 (8-3 mice died=5) |
| n3D | |
| 16 weeks | 8 |
| 1 year | 7 (8-1 mouse died=7) |
| 2 years | 5 (8-3 mice died=5) |
| WD | |
| 16 weeks | 8 |
| 1 year | 7 (8-1 mouse died=7) |
| 2 years | 3 (8-5 mice died=3) |
| | P value |
| 2-way ANOVA | |
| Age | <0.001*** |
| Diet | <0.001*** |
| Tukey's multiple comparison | |
| 16 weeks | |
| LFD vs n3D | 0.550 |
| LFD vs WD | 0.006** |
| n3D vs WD | 0.087 |
| 1 year | |
| LFD vs n3D | 0.171 |
| LFD vs WD | 0.035 |
| n3D vs WD | 0.752 |
| 2 years | |
| LFD vs n3D | 0.489 |
| LFD vs WD | 0.011 |
| n3D vs WD | <0.001*** |
| LFD | |
| 16 weeks vs 1 year | 0.054 |
| 16 weeks vs 2 years | 0.802 |
| 1 year vs 2 years | 0.024* |
| n3D | |
| 16 weeks vs 1 year | 0.009** |
| 16 weeks vs 2 years | 0.018* |
| 1 year vs 2 years | <0.001*** |
| WD | |
| 16 weeks vs. 1 year | 0.193 |
| 16 weeks vs 2 years | 0.938 |
| 1 year vs 2 years | 0.589 |

Panel F (Male sWAT TG uptake)

| | # of animals |
|---|---|
| LFD | |
| 16 weeks | 8 |
| 1 year | 8 |
| 2 years | 5 (8-3 mice died=5) |
| n3D | |
| 16 weeks | 8 |
| 1 year | 7 (8-1 mouse died=7) |
| 2 years | 5 (8-3 mice died=5) |
| WD | |
| 16 weeks | 8 |
| 1 year | 7 (8-1 mouse died=7) |
| 2 years | 3 (8-5 mice died=3) |
| | P value |
| 2-way ANOVA | |
| Age | <0.001*** |
| Diet | 0.008** |
| Tukey's multiple comparison | |
| 16 weeks | |
| LFD vs n3D | 0.615 |
| LFD vs WD | 0.585 |
| n3D vs WD | 0.139 |
| 1 year | |
| LFD vs n3D | 0.461 |
| LFD vs WD | 0.297 |
| n3D vs WD | 0.952 |
| 2 years | |
| LFD vs n3D | 0.030* |
| LFD vs WD | 0.020* |
| n3D vs WD | 0.862 |
| LFD | |
| 16 weeks vs 1 year | 0.550 |
| 16 weeks vs 2 years | 0.011* |
| 1 year vs 2 years | <0.001*** |
| n3D | |
| 16 weeks vs 1 year | 0.008** |
| 16 weeks vs 2 years | 0.754 |
| 1 year vs 2 years | 0.109 |
| WD | |
| 16 weeks vs. 1 year | 0.272 |
| 16 weeks vs 2 years | 0.962 |
| 1 year vs 2 years | 0.334 |

Panel G (Male BAT TG uptake)

| | # of animals |
|---|---|
| LFD | |
| 16 weeks | 8 |
| 1 year | 8 |
| 2 years | 5 (8-3 mice died=5) |
| n3D | |
| 16 weeks | 8 |
| 1 year | 7 (8-1 mouse died=7) |

| | |
|---|---|
| 2 years | 5 (8-3 mice died=5) |
| WD | |
| 16 weeks | 8 |
| 1 year | 7 (8-1 mouse died=7) |
| 2 years | 3 (8-5 mice died=3) |
| | P value |
| 2-way ANOVA | |
| Age | <0.001*** |
| Diet | 0.460 |
| Tukey's multiple comparison | |
| 16 weeks | |
| LFD vs n3D | 0.094 |
| LFD vs WD | 0.168 |
| n3D vs WD | <0.001*** |
| 1 year | |
| LFD vs n3D | 0.714 |
| LFD vs WD | 0.366 |
| n3D vs WD | 0.104 |
| 2 years | |
| LFD vs n3D | 0.512 |
| LFD vs WD | 0.284 |
| n3D vs WD | 0.041* |
| LFD | |
| 16 weeks vs 1 year | 0.096 |
| 16 weeks vs 2 years | 0.027* |
| 1 year vs 2 years | <0.001*** |
| n3D | |
| 16 weeks vs 1 year | 0.005** |
| 16 weeks vs 2 years | 0.901 |
| 1 year vs 2 years | 0.040* |
| WD | |
| 16 weeks vs. 1 year | 0.239 |
| 16 weeks vs 2 years | <0.001*** |
| 1 year vs 2 years | <0.001*** |

**Supplementary Figure 8 statistics**

Panel A (Female GTT 2 years)

| | # of animals |
|---|---|
| LFD | 6 (10-4 mice died=6) |
| n3D | 4 (10-6 mice died=4) |
| WD | 2 (10-8 mice died=2) |

Panel B (Male GTT 2 years)

| | # of animals |
|---|---|
| LFD | 6 (8-2 mice died=6) |
| n3D | 5 (8-3 mice died=5) |
| WD | 3 (8-5 mice died=3) |

Panel C (female insulin 2 years)

| | # of animals |
|---|---|
| LFD | 6 (10-4 mice died=6) |
| n3D | 4 (10-6 mice died=4) |
| WD | 2 (10-8 mice died=2) |

**Supplementary Figure 9 statistics**

Panel A (Female ITT 2 years)

| | # of animals |
|---|---|
| LFD | 6 (10-4 mice died=6) |
| n3D | 4 (10-6 mice died=4) |
| WD | 2 (10-8 mice died=2) |

Panel B (Male ITT 2 years)

| | # of animals |
|---|---|
| LFD | 6 (8-2 mice died=6) |
| n3D | 5 (8-3 mice died=5) |
| WD | 3 (8-5 mice died=3) |
